# pastForward: a Snakemake pipeline for ancient and historical DNA with eukaryote-wide taxonomic screening and tracking of copy-number variation

**DOI:** 10.64898/2026.08.07.743613

**Authors:** Sarah Saadain, Martin Kapun, Robert Kofler

**Affiliations:** Department Biological Sciences and Pathobiology, University of Veterinary Medicine, Vienna, Austria; Vienna Graduate School of Population Genetics, Vetmeduni Vienna, Vienna, Austria; Central Research Laboratories, Natural History Museum Vienna, Vienna, Austria

## Abstract

Ancient and historical DNA has the potential to resolve many open questions in biology. While pipelines for processing ancient and historical DNA exist, none combine user-friendly, configurable processing with copy number variation tracking and targeted taxonomic profiling. Therefore, we developed pastForward, a fully automated Snakemake pipeline that integrates all analysis steps from raw reads to damage-rescaled BAM files in a single reproducible workflow. It performs ancient and historical DNA processing, including adapter trimming, read merging, deduplication, damage assessment, quality rescaling, and generates interactive reports summarizing the endogenous read content, library complexity, and breadth and depth coverage statistics. These reports allow users to rapidly assess the quality of sequencing data. It handles single- and paired-end NGS libraries. Mapping to multiple reference sequences is supported, facilitating co-analysis of host and endosymbiont sequences and genotyping of marker genes such as COI.

pastForward further integrates two novel tools. ECMSD (Efficient Comprehensive Mitochondrial Sequence Detector) screens each library for eukaryotic DNA by aligning reads against a mitochondrial reference database. The presence of bacteria, archaea and viruses is detected in parallel with Centrifuge. REVEAL (Read-based Estimation and visualization of Element Abundance and Loci) quantifies and visualizes copy number variation of genetic features, such as transposable elements (TEs) or gene duplications.

Two case studies demonstrate the usage of the pipeline. Using pastForward on dog genomic time series, including Neolithic samples, we confirm that the copy number of *AMY2B*, which encodes the starch-digesting enzyme amylase, increased during domestication. From historical *D. melanogaster* genomes, we recover the recent invasion of the transposable element *opus*. It is absent in specimens from the 1800s and present from 1933 onward. By efficiently processing large numbers of samples, pastForward facilitates longitudinal tracking of genomic features in diverse species.

## Introduction

Archaeological remains and historical museum collections often preserve genetic material that offers direct windows into past populations, biodiversity, and evolutionary changes over time [Orlando et al., 2012]. The sequencing of ancient (aDNA) and historical DNA (hDNA) has transformed fields ranging from human prehistory [Green et al., 2010] to conservation genomics [Murray et al., 2017]. It enables researchers to address important open questions that cannot be answered from contemporary samples alone. These include, for example, the proportion of archaic admixture in human evolution [Green et al., 2010] and the genomic history of domestication [Bergström et al., 2020]. The emerging field of museomics has leveraged natural history collections to track evolutionary dynamics directly across time and space [Davis and Knapp, 2025, Kapun et al., 2025].

Recent technological and methodological advances have accelerated both the scale and taxonomic breadth of aDNA and hDNA projects. These include DNA extraction from bones [Pinhasi et al., 2019] and minimally invasive extraction from dry-pinned insects in museum collections [Gilbert et al., 2007, Korlević et al., 2021]. They also include single-stranded library preparation protocols [Kapp et al., 2021], DNA repair methods [Jónsson et al., 2013, Skoglund et al., 2014], and decreasing sequencing costs. Besides whole-genome sequencing, amplicon sequencing of marker genes remains widely used for species identification and phylogenetic placement of ancient specimens, especially where reference genomes are unavailable [Thomsen et al., 2009]. Mitochondrial markers, such as cytochrome c oxidase subunit I (COI), are the most common choice.

Across whole-genome and marker gene sequencing, bioinformatic data processing remains computationally demanding. Post-mortem degradation leaves the DNA heavily fragmented, with typical aDNA inserts ranging from 35 to 100 bp [Hofreiter et al., 2001, Lindahl, 1993]. Such short fragments require dedicated read merging, trimming, and mapping strategies [Prüfer et al., 2010]. Contamination may further complicate interpretation. It can stem from exogenous DNA of other organisms or from modern DNA of the target species [Skoglund et al., 2014]. Depurination leads to DNA nicking and fragmentation. Deamination, particularly at single-stranded ends of fragments, additionally results in C*→*T and G*→*A substitutions [Briggs et al., 2007, Sawyer et al., 2012]. These mutations substantially impact genomic analyses, as they impair alignment of reads and bias variant calling. To avoid these problems it is thus necessary to perform damage profiling and base-quality rescaling [Jónsson et al., 2013].

Standard pipelines designed for modern genomics assume long and undamaged reads. They are therefore poorly suited for aDNA/hDNA data. Default quality thresholds discard authentic but short reads, and the default k-mer lengths in metagenomic classifiers often do not work well at these fragment lengths [Wood et al., 2019]. Suitable pipelines must address each of these problems. They should also merge and trim short fragments, screen for endogenous and exogenous DNA, and remove PCR duplicates, which may be abundant in some samples. For whole-genome data, they should additionally rescale base quality scores at read ends to reflect deamination patterns. They should provide comprehensive summary statistics, which allow researchers to assess sample quality and decide whether additional sequencing is necessary. These requirements demand a multi-step workflow that many genomics pipelines are not designed for.

Moreover, ancient and historical DNA sequencing projects are often highly specialized and tailored to the individual specimen, sample type, and research question. Pipelines should thus be flexible and configurable to accommodate these project-specific requirements.

A growing number of specialized bioinformatic tools address individual steps of the aDNA/hDNA workflow. The program fastp [Chen et al., 2018] allows merging overlapping sequence pairs and trimming adapter sequences from reads, even when their exact adapter sequence is unknown. Short reads (typical for aDNA/hDNA datasets) can be reliably mapped with BWA-aln [Li and Durbin, 2009], BWA-mem2 [Vasimuddin et al., 2019] or minimap2 [Li, 2018]. mapDamage2 [Jónsson et al., 2013] identifies post-mortem damage patterns and provides rescaled base qualities if a reference genome is available. Library-aware duplicate removal is performed by DeDup [Peltzer et al., 2016] and the taxonomic composition of a library can be screened with Kraken2 [Lu et al., 2022] or MALT [Herbig et al., 2016].

Established pipelines such as PALEOMIX [Schubert et al., 2014], nf-core/eager [Yates et al., 2021], and mapache [Neuenschwander et al., 2023] assemble subsets of these tools into automated workflows. Four gaps nevertheless remain (Table 3). First, taxonomic screening is either absent or leaves the choice of the reference database to the user. The pre-built indices of k-mer based classifiers must be held in memory in full at every invocation, which exceeds the memory of a typical workstation [Lu et al., 2022]. Memory-capped builds and custom indices offer a smaller footprint, but they classify fewer reads or require substantial memory to build. Second, when testing for contamination with exogenous DNA using mtDNA sequences, nuclear mitochondrial DNA segments (NUMTs) can produce false-positive contamination signals in many pipelines [Hebsgaard et al., 2005]. Third, testing for copy number variation of features across samples is not supported by current pipelines, despite its importance for tracking changes in gene copy number over time [Botigué et al., 2017] and identifying transposable element invasions [Scarpa et al., 2024]. Finally, several existing pipelines depend on workflow managers that require extensive prior knowledge.

To address these issues, we developed pastForward, a Snakemake based [Mölder et al., 2021] bioinformatics pipeline that integrates the complete aDNA/hDNA processing workflow. The user provides raw read files and one or multiple reference sequences. The pipeline performs read processing (adapter trimming, filtering of low quality reads), taxonomic screening, mapping, deduplication, damage profiling, base-quality rescaling, and breadth and depth coverage quantification. The pipeline produces analysis-ready BAM files and accompanying quality reports.

Two novel components are integrated into pastForward: (1) ECMSD determines the taxonomic composition of a library by aligning reads to a curated NCBI RefSeq database of mitochondrial genomes. It distinguishes authentic mitochondrial signals from NUMT artifacts based on breadth coverage; and (2) REVEAL which estimates and visualises the copy number of features of interest (e.g. transposons, genes, etc.) across multiple samples. This enables temporal tracking of the abundance of these features in ancient, historical, and modern datasets.

pastForward offers a reproducible, quick-to-install, modular, and easily configurable solution for processing and evaluating large-scale historical and ancient genomics datasets.

## Materials and Methods

### Overview

pastForward is organized into four modules (Fig. 1), each of which can be independently enabled or disabled. Default parameter values tailored to aDNA/hDNA analyses are provided in a config-file. All parameters can be adjusted.

**Figure 1:**
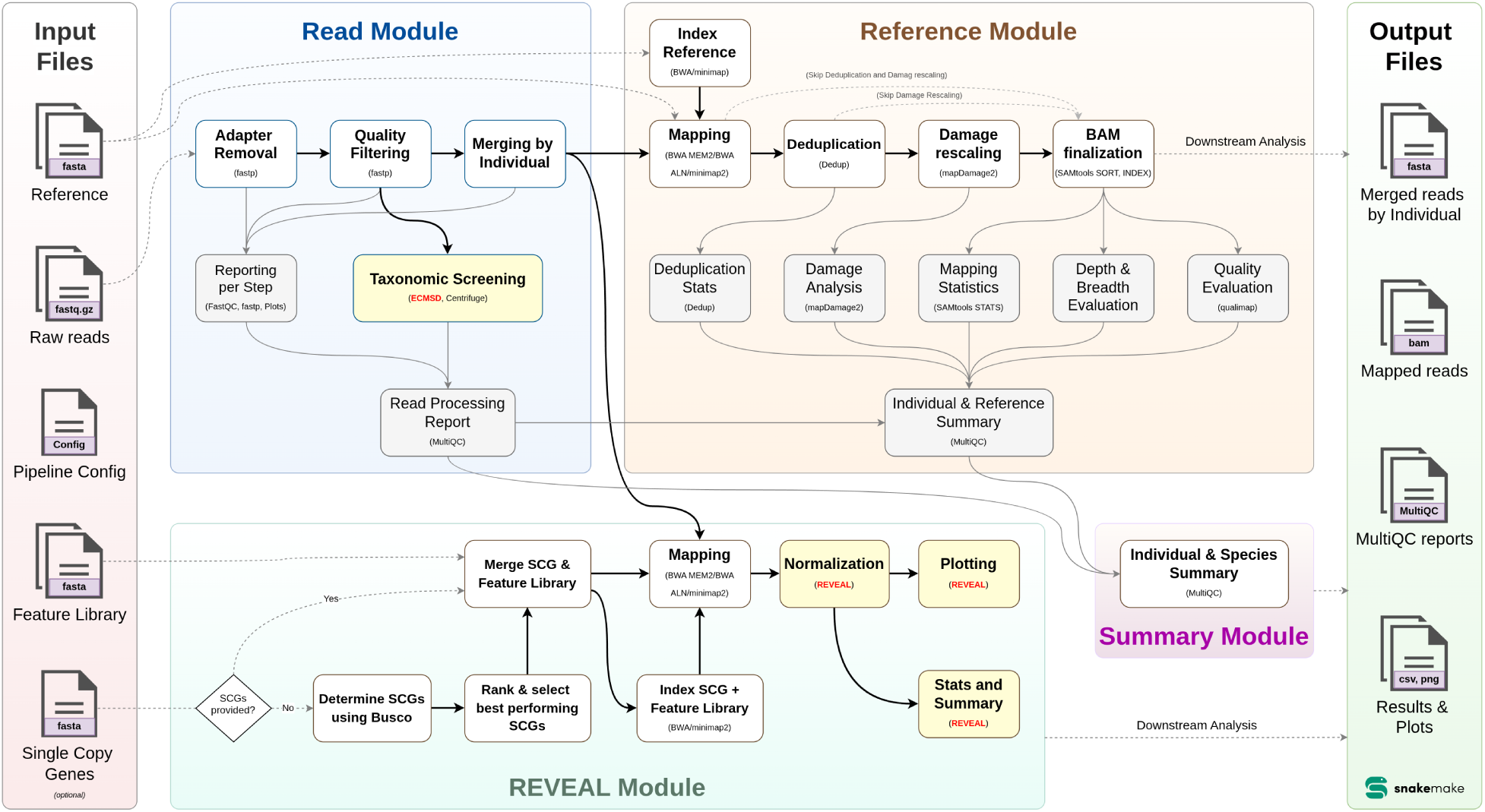
Overview of the pastForward pipeline. The workflow is divided into four modules: the Read Module covers staging, adapter trimming, merging of overlapping pairs, quality filtering, and taxonomic screening; the Reference Module performs read mapping, duplicate removal, damage profiling, base-quality rescaling, and coverage quantification. The Summary module consolidates the output of the Reads- and Reference Modules into a single html MultiQC report. The optional REVEAL Module quantifies and visualises the abundance of user-defined genomic features. Dashed arrows indicate optional steps; solid arrows indicate required steps.

Snakemake tracks file dependencies and executes only the steps needed to create the desired output files. An interrupted run therefore resumes at the missing steps, and a step is repeated only when its input files have changed. Intermediate files are removed automatically when they are no longer needed. Snakemake further manages the parallelization of multi-threaded jobs.

Apart from installing Snakemake itself, all software dependencies are managed by pastForward via the conda environments; hence, no additional manual installation of tools is needed.

pastForward requires raw, compressed sequencing reads in FASTQ format (.fastq.gz) and reference sequences in FASTA format (e.g., genomes, markers) as input. When using REVEAL, genomic feature sequences, such as transposons or genes, must additionally be provided as FASTA files. A set of single-copy genes (SCGs) may be provided in the same format. If none are given, pastForward identifies them automatically.

To run pastForward, users define one or more focal species with reference genomes and/or marker genes in the configuration file. pastForward uses this information to determine which input files should be processed. Per species, the user defines a prefix (e.g., Dmel, Dsim) and provides the input files in a folder with the same name, thus associating the reads with the corresponding reference(s). pastForward supports concurrent processing of multiple species, multiple references per species, as well as multiple samples per species. The output is organized into four categories, each corresponding to a module.

The **Read Module** generates trimmed and filtered read files (.fastq.gz) per sample and screens them taxonomically with ECMSD and Centrifuge [Kim et al., 2016].

The **Reference Module** aligns these reads and generates deduplicated and damage rescaled BAM files for each sample and reference, together with quality reports and summary statistics.

The **Summary Module** aggregates the results of both modules into interactive MultiQC [Ewels et al., 2016] reports.

The **REVEAL Module** estimates and visualises the copy number of genomic features of interest.

### Read Module

The Read Module prepares fastq reads for all downstream steps. Single- and paired-end reads are recognized and processed accordingly. First, adapters are removed with fastp [Chen et al., 2018] where the adapter sequence is automatically detected (unless provided by the user). Poly-X tails are trimmed (trim poly x), and reads exceeding a threshold number of ambiguous bases (n base limit) or unqualified bases (unqualified percent limit) are discarded. fastp also merges paired-end reads when possible. Merged and unmerged reads are joined into a single file and passed through quality filtering, which removes reads that are too short (min length) or of insufficient base quality (min quality). Each library is then screened taxonomically, as described in the next paragraph. Finally, all libraries from the same sample are concatenated into a single FASTQ file (a sample may have been sequenced multiple times). Samples are identified based on a user specified prefix in the filename.

Screening each library separately shows whether endogenous DNA of the target species is present. It also detects exogenous DNA from other organisms. Low-quality or contaminated libraries can therefore be excluded before further use. Two complementary tools are provided: (1) ECMSD, which identifies eukaryotic taxa and is described in the next subsection, and (2) Centrifuge [Kim et al., 2016], which identifies bacteria, archaea and viruses.

We selected Centrifuge over Kraken2 for two reasons. First, its memory footprint is much smaller. Centrifuge compresses the redundancy among closely related genomes, so its index stays compact. The pre-built index for bacteria, archaea, viruses and human occupies 7.9 GB. The corresponding Kraken2 index occupies 233 GB and covers the same groups plus protozoa, fungi and plants [Lu et al., 2022]. Second, Centrifuge classifies shorter fragments. Kraken2 requires reads of at least the k-mer length, 35 bp by default, and a shorter fragment yields no k-mers at all [Wood et al., 2019]. Centrifuge instead scores exact matches of variable length, from a minimum of 22 bp [Kim et al., 2016]. In our historical *D. melanogaster* libraries about 10 % of quality-filtered reads are shorter than 35 bp and are therefore invisible to Kraken2.

pastForward downloads and configures the Centrifuge index automatically. Human is omitted from the Centrifuge summary, because human DNA is already covered by the eukaryotic screening of ECMSD.

### ECMSD: Efficient Comprehensive Mitochondrial Sequence Detector

ECMSD is a lightweight tool for taxonomic screening. It aligns reads against a curated database of complete mitochondrial genomes. It therefore classifies eukaryotes, including metazoans.

Metazoans are the gap left by general-purpose metagenomic classifiers. Their pre-built databases cover bacteria, archaea and viruses, and some also fungi, protozoa and plants. No metazoan is included other than human [Wood et al., 2019, Lu et al., 2022, Kim et al., 2016]. Screening for further metazoans requires a custom database, which is computationally demanding to build.

Mitochondrial genomes offer a viable compromise between taxonomic breadth and database size. Curated reference sequences are available for more than 10,000 taxa. The trade-off is that only the mitochondrial genome is screened. Accuracy may therefore be lower than for tools that evaluate entire genomes. Centrifuge and ECMSD together therefore span the full taxonomic range at low memory cost: prokaryotes and viruses on one side, eukaryotes including metazoans on the other.

ECMSD builds the database automatically during the first run. Curated reference records are downloaded from NCBI RefSeq, which is non-redundant and provides one record per organism. Low-complexity regions are soft-masked with BBMask [Bushnell, 2014]. Each reference is then linked to its NCBI taxonomy identifier, which resolves it to species level and adds genus, family, order, class and phylum. The finished database occupies 0.67 GB on disk.

Reads are aligned to this database with minimap2 [Li, 2018], using the short-read preset (-x sr), and filtered by mapping quality (default: MAPQ *≥* 20). Alignment-based classification is more accurate than alignment-free classification, which suffers from higher false-positive rates, but it is computationally more demanding [Szymanski et al., 2025]. Restricting the reference to mitochondrial genomes keeps this cost low. Analyses based on mitochondria are affected by nuclear mitochondrial insertions (NUMTs). NUMTs produce reads that are mitochondrial in sequence but nuclear in origin, and they have repeatedly caused erroneous taxonomic assignments in aDNA studies [Hebsgaard et al., 2005, Reiss, 2006, Cooper and Poinar, 2000]. A NUMT covers only part of the mitochondrial genome, whereas authentic mitochondrial DNA covers it broadly. ECMSD therefore computes the percentage of each reference that is covered by at least one read, and retains only references above a breadth coverage threshold (default: 25 %). The same filter also removes spurious hits caused by mapping artifacts.

ECMSD reports the taxonomic composition of the mitochondrial read fraction as a table and three plots. These show the read-length distribution per taxon at any taxonomic level, the proportion of classified reads per taxon, and the breadth and depth coverage for each top-ranked reference. The length distribution helps to distinguish genuine aDNA/hDNA from contemporary contamination. Degraded DNA yields an excess of short fragments, while reads from a modern contaminant are longer.

### Reference Module

The Reference Module maps the reads (joint libraries per sample; see above) to one or more reference sequence(s) using BWA-MEM2 [Vasimuddin et al., 2019] (default). BWA-aln [Li and Durbin, 2009] or minimap2 [Li, 2018] may be used as alternatives. The alignment to multiple reference genomes may, for example, facilitate analyzing host and endosymbiont genomes, exploring the suitability of multiple available reference genomes, and the use of marker genes (e.g. COI) in parallel with whole genome analysis.

Following read alignment, PCR duplicates are removed using a modified version of DeDup [Peltzer et al., 2016]. We implemented an improved version with unchanged functionality, but with various performance enhancements, including parallel processing of chromosomes. For a BAM file with 100 million reads, our fork is about 10*×* faster than the original (8 threads). Algorithmic changes, correctness verification, and scaling benchmarks are described in supplementary note 1. After that, pastForward employs mapDamage2 [Jónsson et al., 2013], which can be used for the authentication of aDNA/hDNA, damage profiling, and base-quality rescaling. PCR deduplication and damage rescaling may be skipped, if specified by the user.

The final BAM file is used to compute various downstream quality metrics. Breadth and depth coverage are calculated with samtools depth [Li et al., 2009] and summarized by custom pastForward scripts. Complementary statistics are obtained from samtools stats and Qualimap [Okonechnikov et al., 2016], including the proportion of the genome with read-depths *>*5*×*, *>*10*×*, and *>*30*×*, as well as the median and mean coverage. Library complexity is estimated with Preseq [Daley and Smith, 2013] based on the BAM-files before deduplication, as otherwise this would lead to artificially elevated estimates of library complexity. The resulting library complexity curves allow for assessing whether additional sequencing of an existing library would increase genome-wide coverage or whether the library is saturated. In the second case, novel genomic information may require new DNA-extractions and library preparation, rather than re-sequencing the already existing library.

### REVEAL Module

The REVEAL Module quantifies the relative abundance of user-defined genomic features, such as TEs or genes, across multiple ancient, historical or modern specimen. It is similar to our tool DeviaTE [Weilguny and Kofler, 2019], but represents a faster re-implementation suitable for analysing many different samples and features simultaneously.

Raw reads are mapped with BWA-MEM2 [Vasimuddin et al., 2019] (BWA-aln [Li and Durbin, 2009] or minimap2 [Li, 2018] are optional) to a genomic sequence of interest, such as the consensus sequences of transposable elements or a gene library. Importantly a set of single copy genes (SCG) is also required for normalizing the depth coverage. If no SCGs were provided by the user, pastForward automatically identifies candidate SCGs using BUSCO gene sets, for which the user must specify the appropriate lineage dataset (e.g. drosophilidae odb12) for each species. By normalising the depth coverage of the target features to the SCG coverage, the copy number of the features can be estimated. For example, if a TE has an average depth of 100 and the SCG an average depth of 25, we can estimate that the TE has about 4 copies per haploid genome. This normalization makes abundances comparable across samples regardless of sequencing depth and endogenous DNA content.

The estimates are reported as .tsv tables. REVEAL also generates visualizations. These let users assess the copy number of a TE or gene and the polymorphism among its copies (e.g. SNPs present in some TE insertions but not in others). Depending on the number of investigated samples or features, these plots can be faceted to facilitate comparisons. For example, when REVEAL is used with genomic time-series, it enables direct observation of transposable element invasions within a species across time (see case study below).

### Summary Module

Finally, the ‘Summary Module’ summarizes outputs from the Read Module and the Reference Module into interactive MultiQC HTML reports. Information from all species, samples and references is aggregated, which enables cross-reference and cross-species comparisons. These reports show read counts (raw, trimmed, filtered), taxonomic results (e.g. exogenous DNA, endosymbionts, endogenous DNA content), and genome-wide breadth and depth coverage statistics. They also report library complexity and damage patterns, which can be used to authenticate aDNA/hDNA [Orlando et al., 2012]. The HTML output also enables downloading specific figures and tables as CSV files for downstream analyzes.

### Pipeline tools and versions

All tools used by pastForward are listed in a table in the supplements (Supplementary Table S1). For commonly used bioinformatics tools, we utilised existing Snakemake wrappers from the Snakemake wrappers repository [Mölder et al., 2021] (Supplementary Table S2). For tools without existing wrappers (DeDup [Peltzer et al., 2016], Centrifuge[Kim et al., 2016]), we implemented direct shell calls within the Snakemake workflow. ECMSD and REVEAL have additionally been made available as stand-alone tools via bioconda, enabling their independent use.

## Results

### Comparison to existing pipelines

pastForward is a general purpose pipeline for processing multiple aDNA/hDNA samples from different species (Fig. 1). It orchestrates well-established analyses tools into an easy-to-use stand-alone pipeline and adds our two novel tools, ECMSD and REVEAL.

Apart from these novel features, pastForward differs from other pipelines, such as PALEOMIX [Schubert et al., 2014], nf-core/eager [Yates et al., 2021], and mapache [Neuenschwander et al., 2023] in several respects (Table 3). First, nf-core/eager relies on Nextflow [Di Tommaso et al., 2017]. This requires familiarity with its channel-based dataflow model and with Groovy, a language rarely used in bioinformatics. PALEOMIX is implemented in Python and built around its own custom workflow manager. By contrast, pastForward is implemented in Snakemake, a widely used and well-documented workflow manager.

Second, pastForward is the only one of these three pipelines that configures its databases fully automatically. This includes the Centrifuge [Kim et al., 2016] index and the ECMSD mitochondrial reference database. The setup burden for new users is therefore much lower. PALEOMIX’s metagenomic protocol, by contrast, requires users to source and curate a genome collection for mapping-based profiling themselves. nf-core/eager integrates MALT- and Kraken2-based classification directly, including MaltExtract [Hübler et al., 2019], but the underlying database must be built by the user independently. Mapache offers no integrated taxonomic screening [Neuenschwander et al., 2023].

In contrast to general-purpose pipelines, several pipelines have been developed for specialized tasks such as metagenomic profiling or contaminant/pathogen detection: aMeta [Pochon et al., 2023] provides taxonomic classification for authenticating ancient pathogen and eukaryotic DNA but does not perform deduplication or damage rescaling; HAYSTAC [Dimopoulos et al., 2022] offers Bayesian, damage-aware species identification via competitive mapping against a multi-species database. quicksand [Szymanski et al., 2025] and schmutzi [Renaud et al., 2015] are designed for screening sedimentary aDNA and estimating human mitochondrial contamination. metaBIT [Louvel et al., 2016] focuses on microbial profiling from shotgun data using metagenomics-specific defaults. nf-core/mag [Krakau et al., 2022] instead assembles and bins metagenomes de novo, and authenticates the assembled contigs with PyDamage [Borry et al., 2021] rather than reads mapped to a reference genome.

Another group of workflows facilitates downstream population-genomic analysis rather than aDNA preprocessing. For example, ATLAS [Link et al., 2017] implements genotype-likelihood methods for ancient samples but expects a pre-processed BAM file as input. GenErode [Kutschera et al., 2022], grenepipe [Czech and Expo-sito Alonso, 2022], and PopGLen [Nolen, 2025] are Snakemake pipelines for assessing genome-erosion, variant calling, and population genomics. However, they lack integrated taxonomic screening.

### Case Study 1: aDNA from neolithic canids reveals increase of *AMY2B* copy numbers during domestication

To demonstrate the utility of pastForward we performed two case studies. First, we evaluated pastForward with aDNA and tested whether our approach is able to reproduce the previously detected copy number variation of *AMY2B* genes in dogs [Axelsson et al., 2013]. We therefore employed a genomic time-series dataset spanning canine domestication, from *∼*7,000-year-old Neolithic dog remains to modern breeds [Botigué et al., 2017]. We additionally included wolf resequencing data as an outgroup, approximating the pre-domestication state of *AMY2B* copy number. Sample accessions and metadata are listed in Supplementary Table S3.

ECMSD classified mitochondrial reads across all samples as canid (Table 1). In the modern wolf (Clup01), *Canis lupus* accounted for 60.9% of classified reads. Its mitogenome was covered to 52.4%, well above the 25% breadth threshold that separates authentic host signal from NUMTs and mapping artifacts. The remaining reads were attributed to closely related canids (*C. rufus*, *C. simensis*, *C. aureus*, *C. lupaster* ). In modern domestic dogs (German shepherds; Clup04, Clup05) 100% of reads (34,605 and 28,542) were assigned to *C. lupus* (56.9% and 64.9% covered). In the Neolithic dogs *C. lupus* accounted for 75.5% and 72.8% of classified reads in Clup03 (7,000y) and Clup02 (5,000y), with breadth coverage of 36.7% and 34.1%. The read composition and coverage statistics for Clup02 are shown in Fig. 2, with full values for all samples given in Table 1. The corresponding ECMSD plots for the remaining samples are provided as Figs. S6–S9.

**Figure 2:**
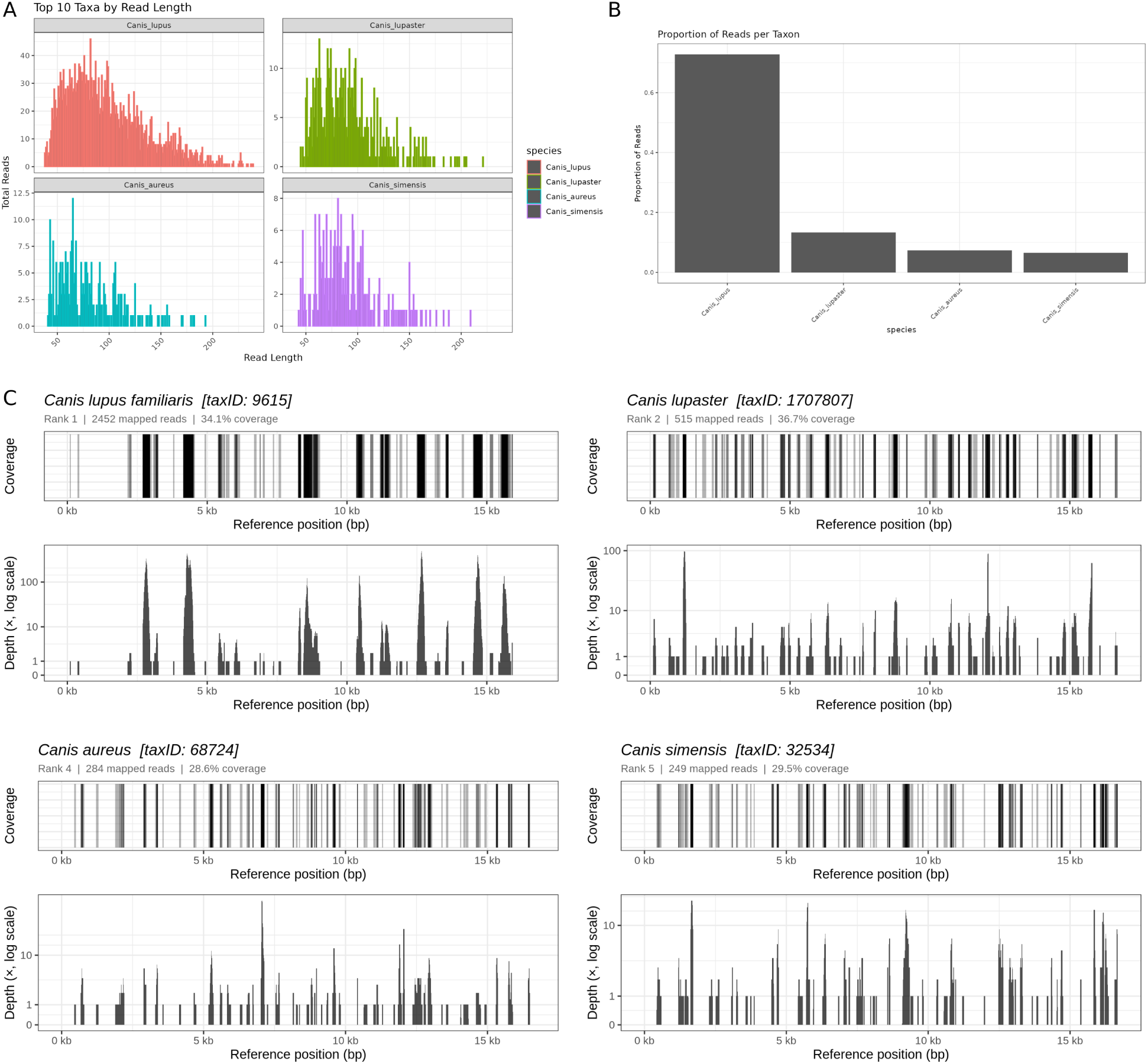
Example of ECMSD output for Clup02, a 5,000y old Neolithic dog [Botigué et al., 2017]. (A) Read-length distribution of the four most abundant taxa. The short fragment lengths seen in all four taxa is typical for degraded aDNA. (B) Proportion of mitochondrial reads assigned to each canid taxon; *Canis lupus* accounts for the majority (72.8% of the classified reads), while the remaining reads were classified as *C. lupaster*, *C. aureus*, and *C. simensis*. (C) Coverage of the *C. lupus* mitochondrial genome (taxID 9615; 2,452 mapped reads): breadth coverage across the genome (top) and per-base depth on a log scale (bottom), showing that reads are distributed across the mitogenome (34.1% breadth).

**Figure 3:**
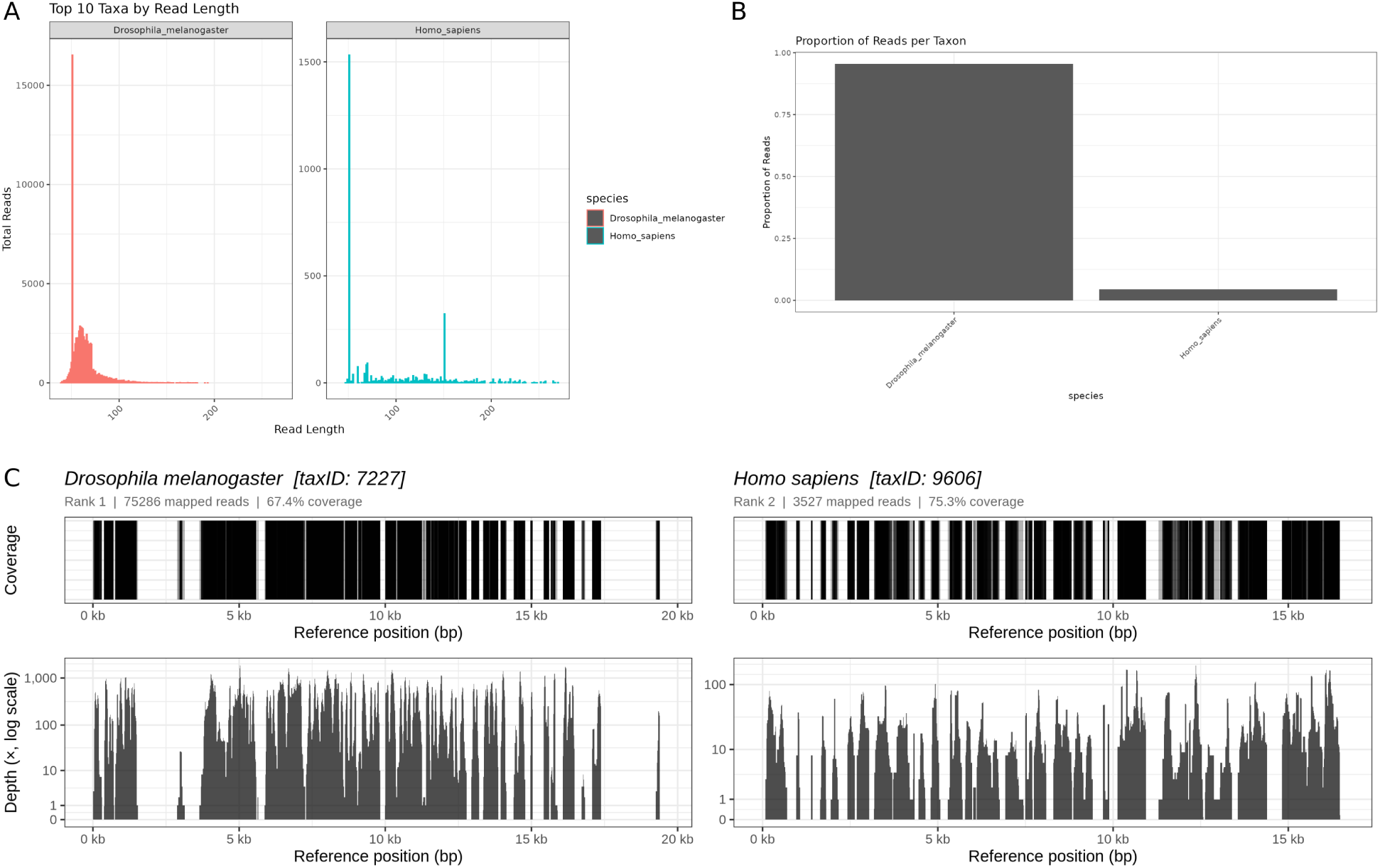
ECMSD output for Dmel1800s-1-SL04, a *∼*200-year-old *D. melanogaster* specimen from a museum collection [Shpak et al., 2023]. (A) Read-length distributions for the two most abundant taxa: *D. melanogaster* and *Homo sapiens*. (B) Proportion of mitochondrial reads per taxon. (C) Breadth and depth coverage of the mitochondrial reference for each taxon. *D. melanogaster* (taxID 7227) ranks first with 75,286 mapped reads and 67.4% of the reference covered; *Homo sapiens* (taxID 9606) ranks second with 3,527 mapped reads and 75.3% covered. Both references exceed the breadth coverage threshold, indicating that the human signal reflects genuine low-level contamination rather than a mapping artifact or NUMT.

**Table 1:** Mitochondrial read composition and coverage for the *Canis lupus* samples Clup01-Clup05.

| Sample | SRA Accession | Input reads | Classified<br>mtDNA reads | Taxon | Reads (%) | Mean depth coverage | mt genome<br>covered (%) |
| --- | --- | --- | --- | --- | --- | --- | --- |
| **Clup01**<br>**modern wolf** | SRR780933 | 155,039,287 | **128,249** | *Canis lupus* | 60.85% | 332.9× | 52.4% |
|  |  |  |  | *Canis rufus* | 21.89% | 139.3× | 29.6% |
|  |  |  |  | *Canis simensis* | 7.13% | 67.5× | 59.4% |
|  |  |  |  | *Canis aureus* | 5.33% | 27.2× | 53.7% |
|  |  |  |  | *Canis lupaster* | 4.46% | 34.3× | 66.9% |
|  |  |  |  | *Canis latrans* | 0.34% | 3.1× | 31.3% |
| **Clup02**<br>**5,000y dog, Neolithic** | SRR3417116 | 524,081,302 | **3,848** | *Canis lupus* | 72.77% | 11.7× | 34.1% |
|  |  |  |  | *Canis lupaster* | 13.38% | 2.1× | 36.7% |
|  |  |  |  | *Canis aureus* | 7.38% | 1.0× | 28.6% |
|  |  |  |  | *Canis simensis* | 6.47% | 0.9× | 29.5% |
| **Clup03**<br>**7,000y dog, Neolithic** | SRR3417117 | 627,760,462 | **6,579** | *Canis lupus* | 75.48% | 18.7× | 36.7% |
|  |  |  |  | *Canis lupaster* | 13.00% | 3.4× | 35.9% |
|  |  |  |  | *Canis aureus* | 5.88% | 1.4× | 36.5% |
|  |  |  |  | *Canis simensis* | 5.64% | 1.5× | 34.9% |
| **Clup04**<br>**modern dog, German Shepherd** | SRR1122359 | 315,949,507 | **34,605** | *Canis lupus* | 100.00% | 207.2× | 56.9% |
| **Clup05**<br>**modern dog, German Shepherd** | SRR13339247 | 397,294,532 | **28,542** | *Canis lupus* | 100.00% | 242.4× | 64.9% |

We then mapped the reads genome-wide to the domestic dog reference (GCF 011100685.1). The Neolithic samples mapped at lower rates than the modern ones (71.3-76.9% vs. 99.7-99.9%; Fig. S2), indicating a lower endogenous DNA content. They also showed strong terminal C*→*T damage (23.8-26.8% vs. *≤*0.3%), as expected for aDNA. Breadth coverage remained high across all five specimen (95.7-99.0% of the genome covered; 6.8-21.3*×* mean depth; Fig. S3), providing sufficient depth for the copy-number analysis below. These per-sample metrics are aggregated by the Summary Module into an interactive MultiQC report (Fig. 5); the full report, with detailed read, mapping, and coverage statistics, is provided as supplementary data S1.

Next, we used REVEAL to estimate and visualise copy-number of *AMY2B* (pancreatic alpha-amylase gene; NC 051810.1:47236426–47243598) during dog domestication (Fig. 4). REVEAL normalises depth coverage to single-copy genes, so all copy numbers are haploid, i.e. half of the diploid counts given in the original studies. Our analyses indicate a single copy of *AMY2B* in modern wolf and neolithic dogs, which corresponds to the two diploid copies reported previously [Botigué et al., 2017]. In contrast, the modern dogs show *∼*10 copies (*∼*20 in the diploid genome), capturing the amplification of *AMY2B* associated with dietary starch adaptation during dog domestication [Axelsson et al., 2013].

**Figure 4:**
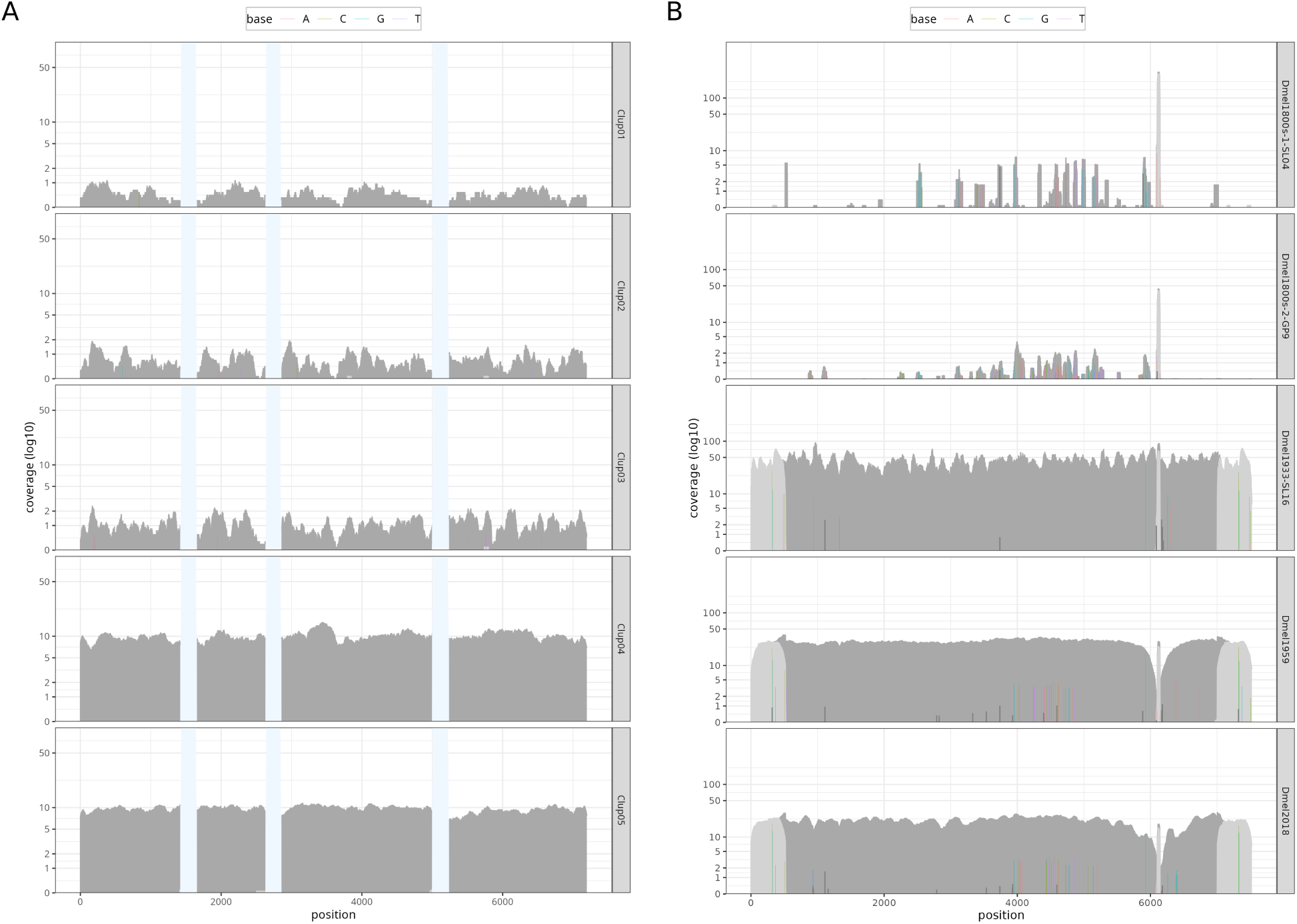
REVEAL plots illustrate copy number variation among samples. (A) Copy number of *AMY2B* gene in wolf (*Canis lupus*; Clup01), two Neolithic dogs (Clup02-03), and modern dog samples (Clup04-05). Copy numbers are haploid, as REVEAL normalises depth coverage to single-copy genes. The samples illustrate an increase of *AMY2B* copy number from wolf to the German Shepherd, consistent with previous work [Axelsson et al., 2013]. Bright blue areas are masked (transposable element fragments). (B) The retrotransposon *opus* is absent in historical *D. melanogaster* samples from the early and mid 1800s but present in later samples, consistent with an invasion of *opus* during the last 2 centuries [Scarpa et al., 2024].

**Figure 5:**
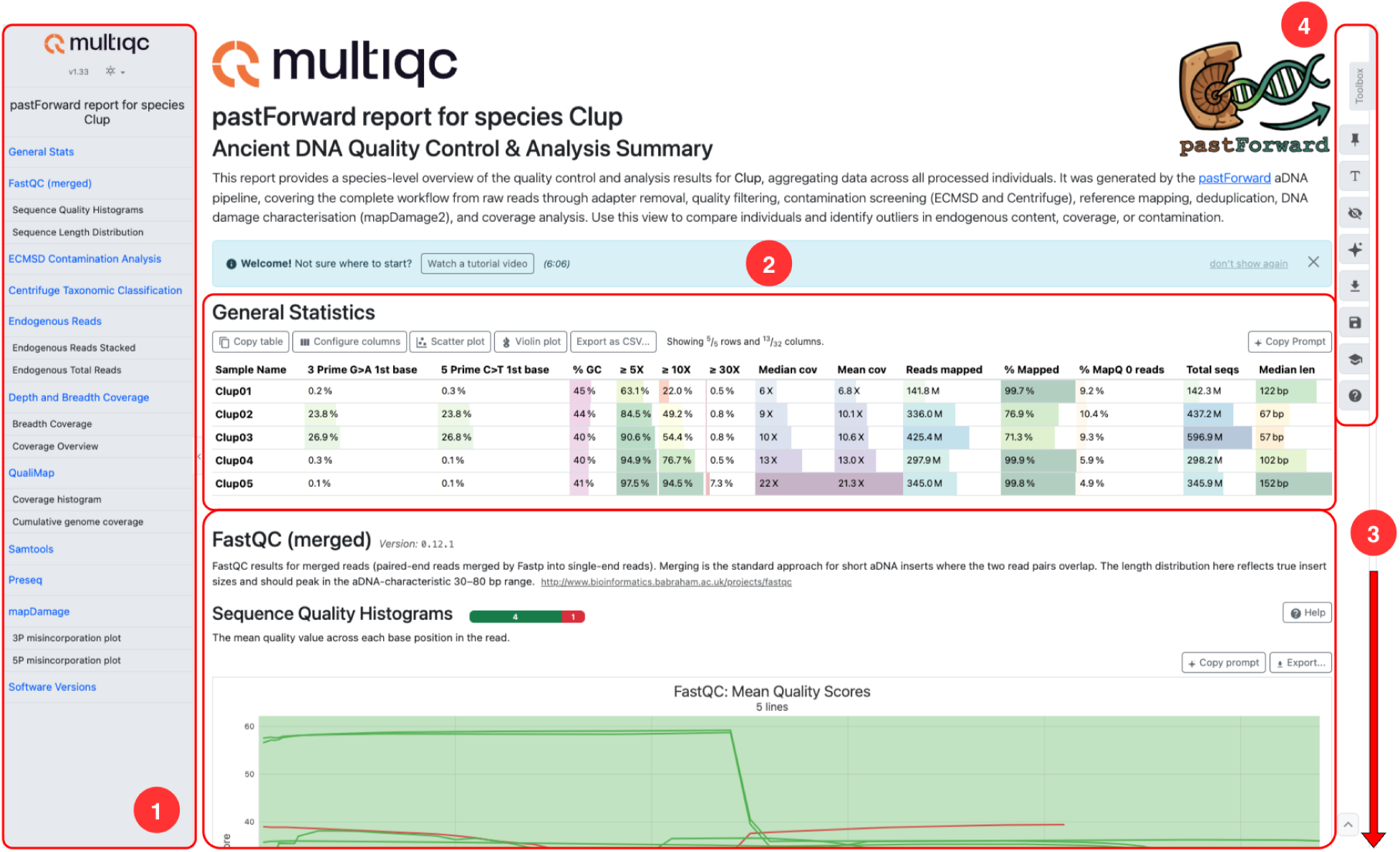
Screenshot of the interactive MultiQC report generated by the Summary Module for the *C. lupus* case study, annotated to highlight its features. (1) The sidebar navigates between the aggregated outputs of the Read and Reference Modules (FastQC, ECMSD taxonomic screening, Centrifuge, endogenous- read quantification, breadth and depth coverage, QualiMap, Samtools, Preseq, and mapDamage2). (2) The General Statistics table provides a quick summary of important per-sample metrics for Clup01-Clup05. (3) Each section presents the corresponding per-sample plots and tables in further detail. Tables and plots can be filtered, sorted, and reconfigured (e.g. switched between table, scatter, and violin views), and the underlying raw data can be exported (e.g. as CSV). (4) A toolbox provides further tools, including AI-assisted summarisation: sections and plots can be exported as ready-made prompts (“Copy Prompt”) for interpretation with external AI assistants. The full report is provided as a PDF in supplementary data S1.

These results demonstrate that pastForward can successfully process a combined dataset of ancient and modern DNA. Moreover, REVEAL successfully reproduces a classical case of copy-number increase of a gene during domestication.

### Case Study 2: hDNA of *D. melanogaster* recreates transposons invasions

We next evaluated pastForward on hDNA in a museomics context. We applied the pipeline to a genomic time series of *D. melanogaster* specimens sampled over the last *∼*200 years, including data from pinned museum specimens [Shpak et al., 2023, Schwarz et al., 2021, Mateo et al., 2018]. In particular, we asked whether we can reproduce the invasion of the retrotransposon *opus*, which Scarpa et al. [2024] first documented in samples from 1933. Pinned insects are among the most abundant, yet least explored specimens in natural history collections. Their relatively recent age and dry preservation contrast with the bone-derived Neolithic material providing a broader evaluation of the pipeline across different samples. Sample accessions and metadata are listed in Supplementary Table S4.

ECMSD identified contamination by humans and *Starmerella* in some samples of the museum specimens. The three historical samples (Dmel1800s-1-SL04, Dmel1800s-2-GP9, and Dmel1933-SL16) have between 95.5-100% *D. melanogaster* reads (Table 2). Dmel1800s-1-SL04 had a small fraction of human reads (4.5%, 17.6*×* mean depth, 75.3% of the human mitogenome covered). Based on the read length distributions, we speculate that human contamination reflects both historical and modern human DNA (Fig. 3). Dmel1933-SL16 carried a small fraction of yeast *Starmerella bacillaris* reads (Fig. S11). Dmel1800s-2-GP9, the modern Dmel2018-STO11 sample, and the laboratory strain from 1959 (Dmel1959) had reads assigned to *D. melanogaster* only, indicating no contamination (Figs. S10, S12 and S13).

**Table 2:** Mitochondrial read composition and coverage statistics for the historical and modern *D. melanogaster* samples, ordered oldest to youngest.

| Sample | SRA Accession | Input reads | Classified<br>mtDNA reads | Taxon | Reads (%) | Mean depth coverage | mt genome<br>covered (%) |
| --- | --- | --- | --- | --- | --- | --- | --- |
| Dmel1800s-1-SL04 | SRR23876564 | 285,478,463 | 78,813 | *D. melanogaster* | 95.52% | 205.5× | 67.4% |
| | | | | *Homo sapiens* | 4.48% | 17.6× | 75.3% |
| Dmel1800s-2-GP9 | SRR23876562 | 383,880,919 | 2,682,688 | *D. melanogaster* | 100.00% | 7282.6× | 83.0% |
| Dmel1933-SL16 | SRR23876579 | 609,508,577 | 311,630 | *D. melanogaster* | 99.53% | 802.9× | 70.9% |
| | | | | *Starmerella bacillaris* | 0.47% | 3.1× | 33.8% |
| Dmel1959 | SRR11846565 | 48,836,898 | 136,770 | *D. melanogaster* | 100.00% | 898.9× | 76.0% |
| Dmel2018-STO11 | SRR5762774 | 40,899,095 | 55,085 | *D. melanogaster* | 100.00% | 252.6× | 66.4% |

**Table 3:** Comparison of pastForward with existing pipelines for ancient and historical DNA analysis. + = supported; *−* = not supported; *∼* = partial support. Taxonomic screening refers to integrated classification of reads against a taxonomic reference in order to determine which taxa are present in a library; *∼* indicates that the classification step exists but requires the user to source or build the reference database. Copy number estimation refers to quantification of the copy number of user-defined genomic features, such as genes or transposable elements, normalised so that estimates are comparable across samples; pipelines that only report per-region sequencing depth are scored *−*. For nf-core/mag, *∼* in the aDNA and damage columns refers to its optional ancient DNA subworkflow (v2.2.0 and later), which estimates damage on assembled contigs rather than on reads mapped to a reference genome.

| Tool | Framework | Pipeline is aDNA specific | Adapter Trimming | Mapping to Reference | Deduplication of BAM | Damage analysis of BAM | Taxonomic screening | Copy number estimation |
| --- | --- | --- | --- | --- | --- | --- | --- | --- |
| nf-core/eager | Nextflow | + | + | + | + | + | ~ | - |
| nf-core/mag | Nextflow | ~ | + | + | - | ~ | ~ | - |
| aMeta | Snakemake | + | + | + | - | - | + | - |
| schmutzi | C++ | + | - | - | - | + | - | - |
| quicksand | Nextflow | + | - | + | + | + | + | - |
| PALEOMIX | Python (custom) | + | + | + | + | + | ~ | - |
| mapache | Snakemake | + | + | + | + | + | - | - |
| ATLAS | C++ | + | - | - | - | + | - | - |
| HOPS/MALT | Java | + | ~ | + | - | + | + | - |
| metaBIT | Bash/Python | - | - | - | - | - | + | - |
| HAYSTAC | Snakemake | + | - | + | - | + | + | - |
| GenErode | Snakemake | ~ | + | + | + | + | - | - |
| grenepipe | Snakemake | - | ~ | + | + | + | - | - |
| PopGlen | Snakemake | ~ | + | + | + | + | - | - |
| **pastForward** | **Snakemake** | + | + | + | + | + | + | + |

Genome-wide mapping to the *D. melanogaster* nuclear reference (GCF 000001215.4) showed high host breadth coverage across all samples, regardless of specimen age (75.6-98.1% of the genome covered; Figs. S4 and S5). Terminal C*→*T damage was much lower than in the bone material (Case Study 1) and declined in younger specimens.

As pastForward may map reads to multiple references in parallel, we also aligned the reads to the *Wolbachia* wMel genome (AE017196.1), jointly assessing host and endosymbiont composition. *Wolbachia* coverage differs strongly among the samples. In the modern sample Dmel2018-STO11, *Wolbachia* is almost absent. In Dmel1959, the *Wolbachia* genome is covered completely (100% breadth, up to 155.8*×* depth). These differences reflect the *Wolbachia* infection status of the samples. The full per-sample MultiQC report, with detailed read, mapping, and coverage statistics, is shown in part in Fig. S1 and provided in full as supplementary data S2.

Using REVEAL, we quantified depth coverage of the *opus* retrotransposon across the *D. melanogaster* time series data (Fig. 4). The two specimens from the 1800s show only sparse, patchy *opus* coverage. Continuous coverage across the whole element first appears in the 1933 specimen and persists in 1959 and 2018. Our time series therefore brackets the *opus* invasion between the mid-1800s and 1933. This recapitulates published observations on the historical invasion of *opus* in *D. melanogaster* populations [Scarpa et al., 2024]. It demonstrates that REVEAL can recover the timing of TE invasions from hDNA in museum collections.

## Discussion

Ancient and historical DNA have moved from a niche into a central focus of evolutionary biology. Natural history and archaeological collections hold time series spanning hundreds to thousands of years, hence evolutionary change can be directly observed rather than indirectly inferred from present-day variation. Such collections also preserve material from extinct taxa and populations that no longer exist. They cover thousands of species and specimens, which makes broad comparative analyses possible without requiring new field sampling [Raxworthy and Smith, 2021, Kapun et al., 2025]. The ongoing biodiversity crisis makes this urgent. For many taxa, collections are the only remaining record of what has already been lost [Davis and Knapp, 2025].

The limiting step often lies in obtaining sufficient amounts of DNA for sequencing and the ensuing data analysis. Museomics brings together archaeologists, taxonomists, curators, ecologists, and evolutionary biologists. Their strength lies in deep domain knowledge, but rarely in bioinformatics. The processing of aDNA/hDNA data remains technically demanding. It requires specialised parameter choices, multiple tools, and careful quality assessment at each stage.

Here we present pastForward, a Snakemake pipeline that integrates the analysis of aDNA/hDNA into a single reproducible workflow. It is applicable to any organism, preservation context, and collection type, from millennia-old bones to century-old pinned insects. It further supports both whole-genome sequencing and marker gene approaches. The broad aim was to enable researchers to focus on the biology rather than on the technical details.

We demonstrate the utility of our novel pipeline with two case studies that process two distinct datasets. These include aDNA from 7,000 and 5,000 year-old dog bones, as well as DNA from dry-pinned insects in museum collections that are up to *∼*200 years old. In both case studies, pastForward recovered biologically meaningful signals, including *AMY2B* copy-number expansion during dog domestication and the historical invasion of the *opus* retrotransposon in *D. melanogaster* populations.

Our novel pipeline also implements two new tools: ECMSD for taxonomic screening and REVEAL for copy number variation estimation. Both extend the functionality of pastForward beyond existing approaches. Taxonomic screening is a critical but challenging step. Existing tools are restricted to human aDNA [Renaud et al., 2015, Weissensteiner et al., 2021], require manually rebuilding databases for short aDNA fragments [Lu et al., 2022], or must hold a whole-genome database in memory [Herbig et al., 2016]. ECMSD avoids these limitations by aligning reads against a compact mitochondrial database. Using ECMSD, we demonstrated that in both our case studies, the reads from the focal species were most abundant. Together with Centrifuge, pastForward thus provides a comprehensive overview of the taxonomic composition of samples.

REVEAL estimates the copy number of user-defined features, such as transposons or genes. As these estimates are comparable across samples, features can be tracked through time series spanning centuries to millennia.

### Limitations and future directions

pastForward does not include probabilistic genotyping tools suited to low-coverage aDNA, such as ANGSD [Korneliussen et al., 2014]. These typically require organism-specific parametrisation and manual tuning by the user. However, we are planning to better link the output of our pipeline to such approaches for follow-up analyses.

The ECMSD mitochondrial reference database is necessarily limited to eukaryotic taxa. Furthermore, it is restricted to curated RefSeq reference records (NC accessions) and excludes RefSeq WGS-derived scaffolds (NW accessions) as these entries frequently represent partial or fragmentary mitochondrial assemblies. Future implementations may include additional curated reference databases to extend the functionality of ECMSD beyond characterization based on mitochondrial sequences only. More generally, the ECMSD database reflects RefSeq completeness at the time of download, so taxonomic groups under-represented in RefSeq will have correspondingly lower classification sensitivity.

The simple installation of pastForward, its automatic database configuration, and Snakemake’s scheduling let pastForward run on a common desktop computer. It therefore fits the infrastructure that museums and medium-sized research groups have. Its scope also extends beyond whole-genome data: marker genes such as COI can also be supplied as references, and the resulting BAM files may be used for phylogenetic inference. Museums can thus analyse their own specimens in-house, from raw reads to phylogenetic placement or the longitudinal tracking of features. Vast numbers of specimens still await sequencing in collections worldwide, and we hope pastForward helps to make these data accessible and to facilitate their analysis.

## Acknowledgments

We thank all members of the Unit genetic and animal breeding for feedback and support. RK, SS, and MK thank the many curators of natural history collections with which we interacted for their interest and support.

## Author Contributions

SS conceived the project and led pipeline development. MK designed and implemented ECMSD. RK and SS designed and implemented REVEAL. SS wrote the manuscript. MK and RK contributed to writing. RK supervised the project.

## Data Availability

pastForward is available at https://github.com/SarahSaadain/PastForward, ECMSD is available at https://github.com/capoony/ECMSD and REVEAL is available at https://github.com/SarahSaadain/REVEAL

