## Supplementary for "pastForward: a Snakemake pipeline for ancient and historical DNA with eukaryote-wide taxonomic screening and tracking of copy-number variation"

### Supplementary Material and Methods

Historical *C. lupus* aDNA libraries were obtained from PRJNA319283 [Botigué et al., 2017]. Modern wolf and dog sequences were retrieved from PRJNA192935, PRJNA233638, and PRJNA685036. For *AMY2B* copy-number analysis, reads were mapped to the *AMY2B* genomic sequence (NC_051810.1:47236426–47243598, *Canis lupus familiaris* ROS_Cfam_1.0). Intronic transposable element remnants were identified using RepeatMasker [Smit et al., 2013-2015] and masked using REVEAL.

Historical *D. melanogaster* sequencing data were obtained from [Shpak et al., 2023] (PRJNA945389). Modern *D. melanogaster* sequences were retrieved from PRJNA634847 and PRJNA329555 [Schwarz et al., 2021, Mateo et al., 2018].

### Supplementary Figures

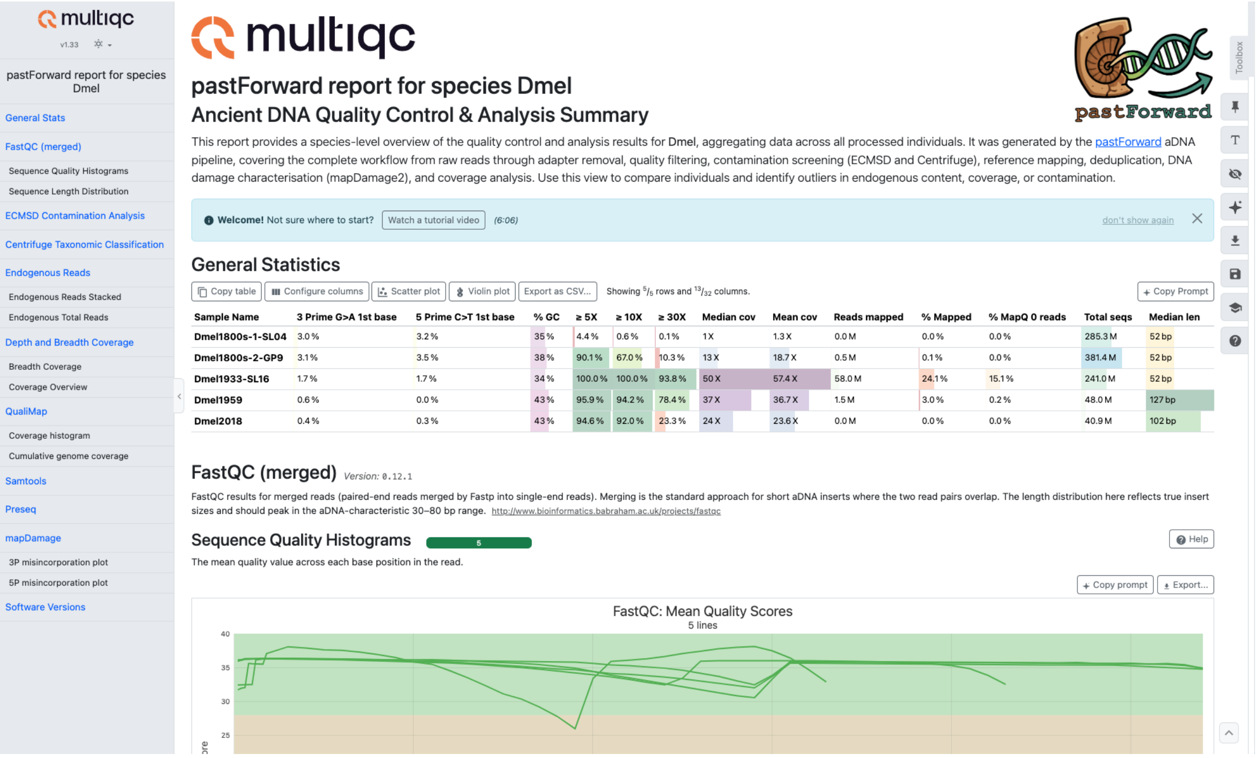


Figure S1: Screenshot of the MultiQC report for the *D. melanogaster* case study. A PDF version of the MultiQC output is available as supplementary data S2.

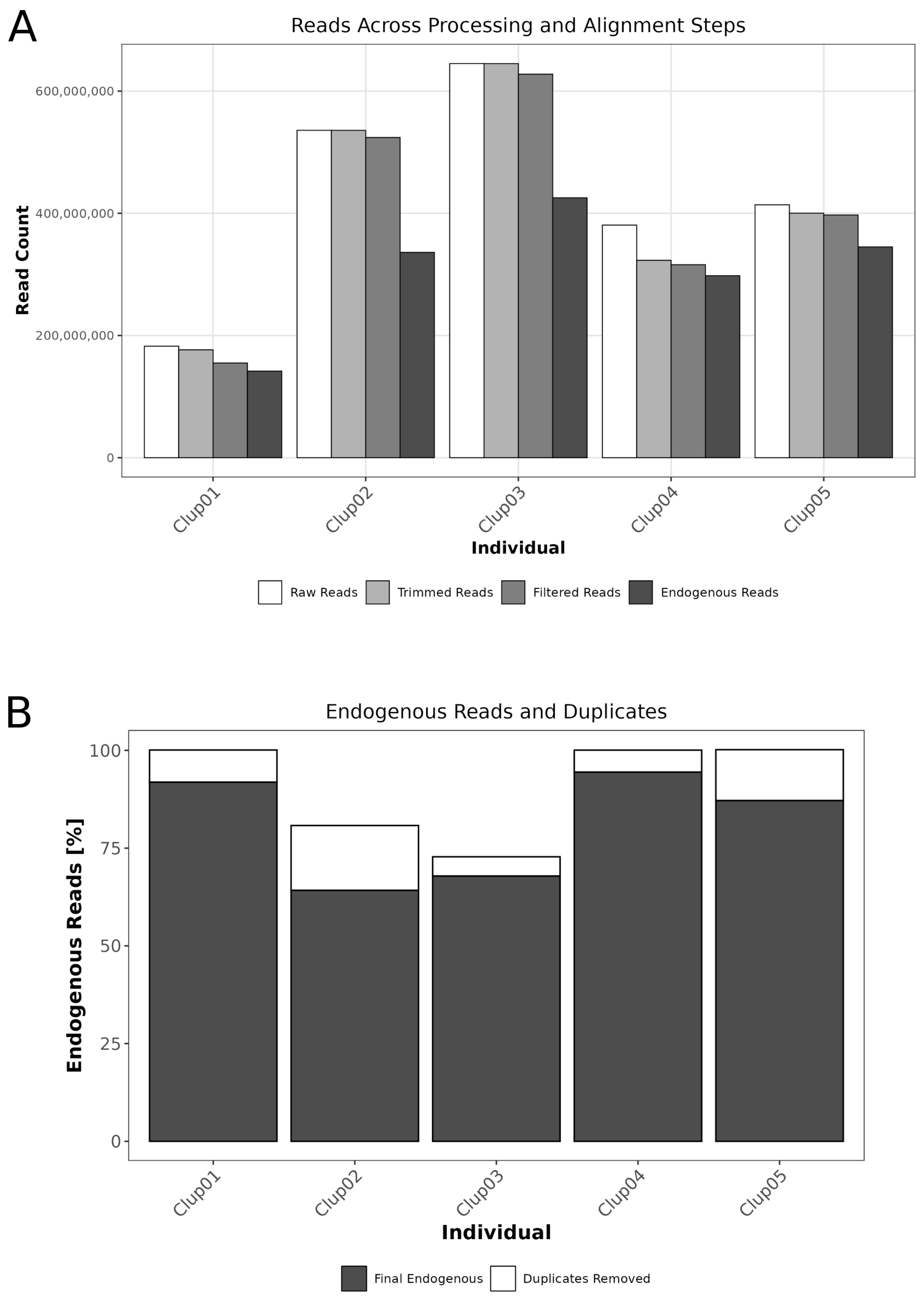


Figure S2: Endogenous *C. lupus* DNA content (percentage of reads mapping to the domestic dog reference, GCF_011100685.1) across the *C. lupus* samples (Clup01–Clup05).

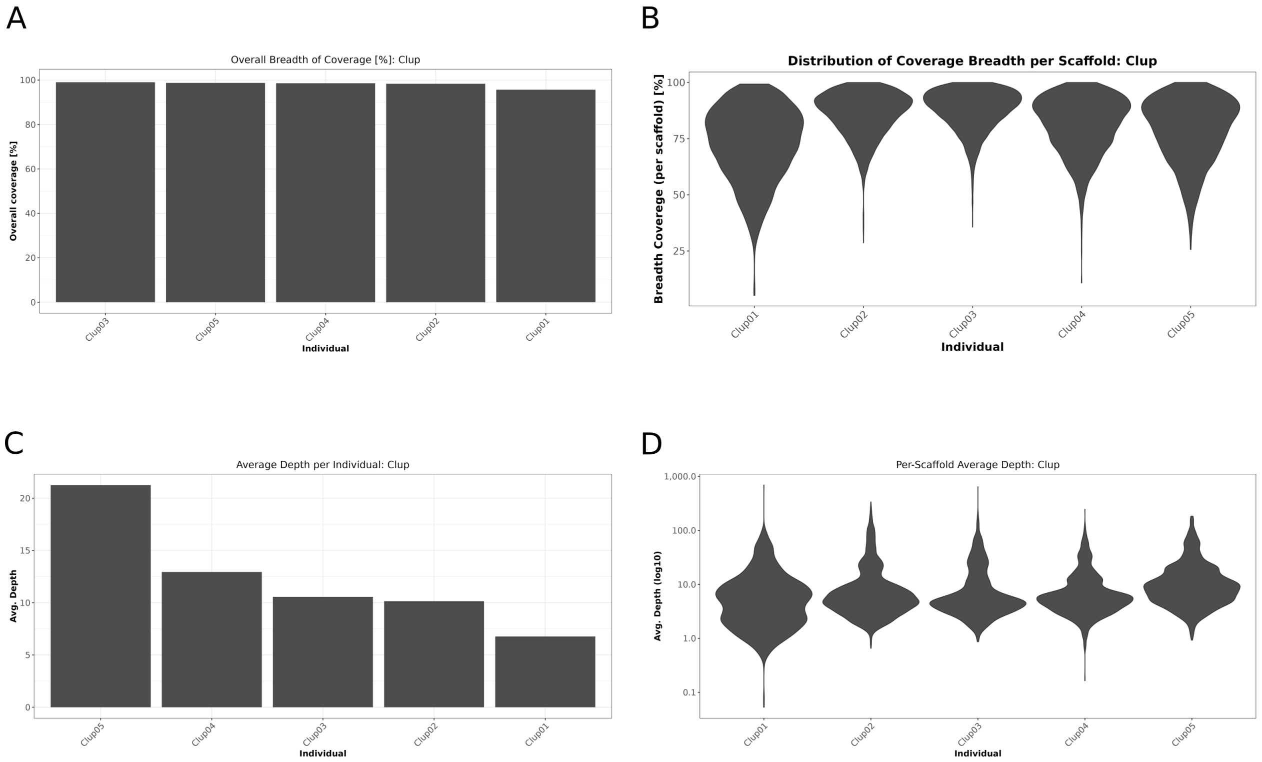


Figure S3: Genome-wide breadth and depth coverage across the *C. lupus* samples following mapping to the domestic dog reference (GCF_011100685.1).

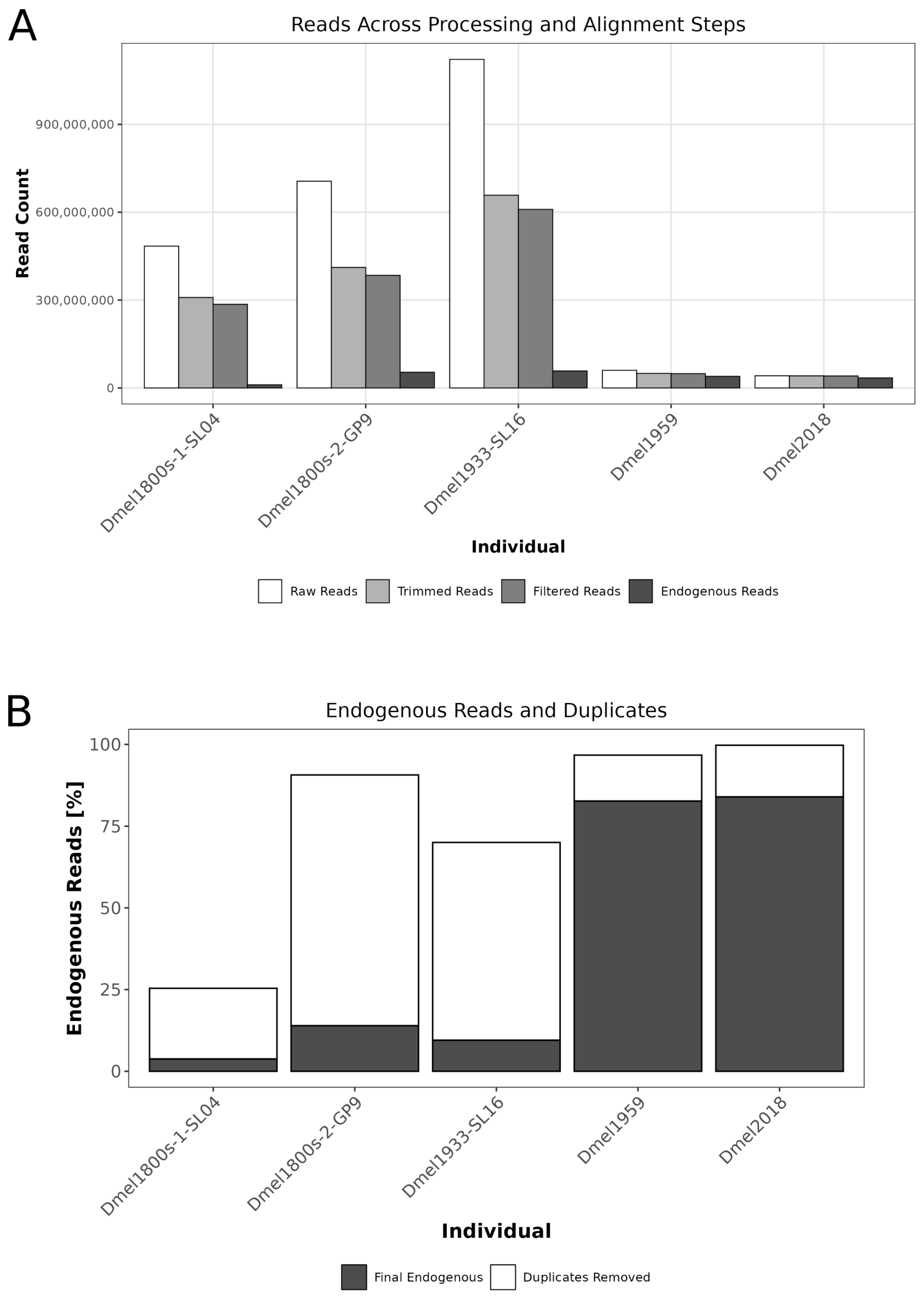


Figure S4: Endogenous *D. melanogaster* DNA content (percentage of reads mapping to the *D. melanogaster* nuclear reference, GCF_000001215.4) across the *D. melanogaster* time-series samples (Dmel1800s-1-SL04 through Dmel2018-STO11).

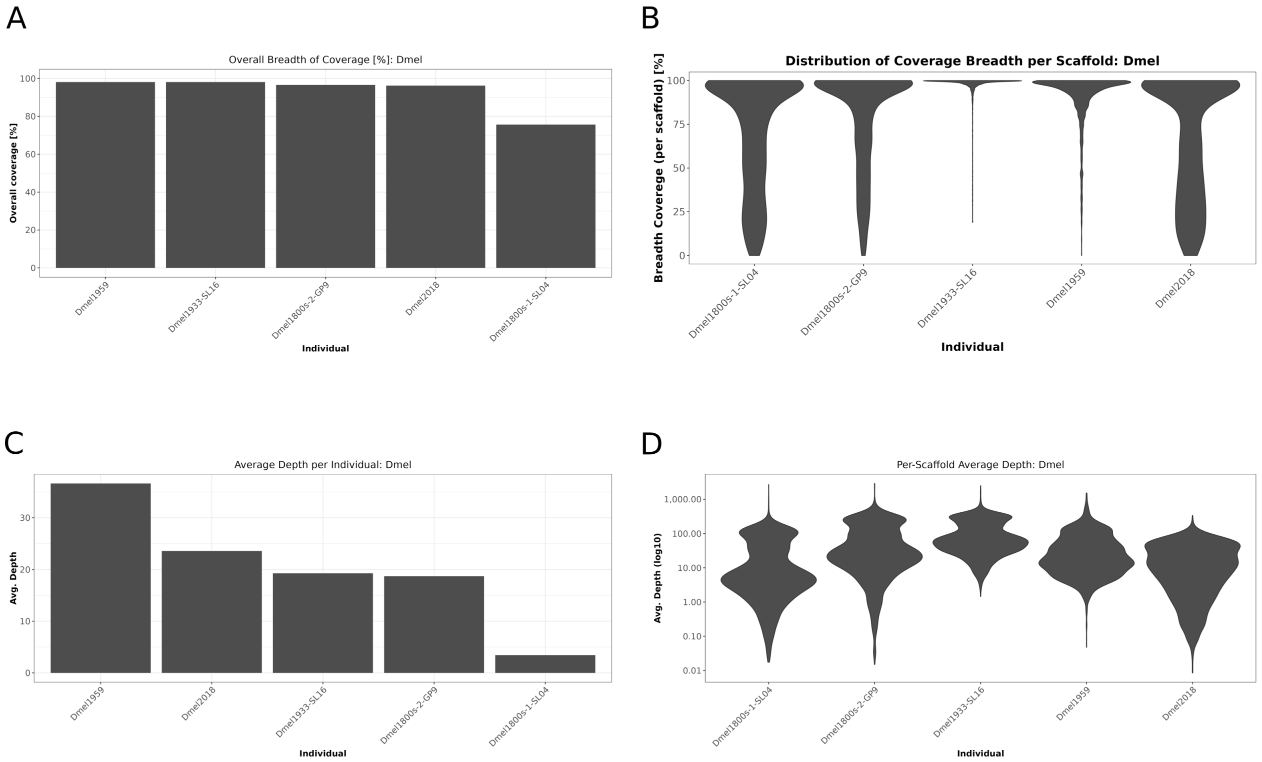


Figure S5: Genome-wide breadth and depth coverage across the *D. melanogaster* time-series samples following mapping to the *D. melanogaster* nuclear reference (GCF_000001215.4).

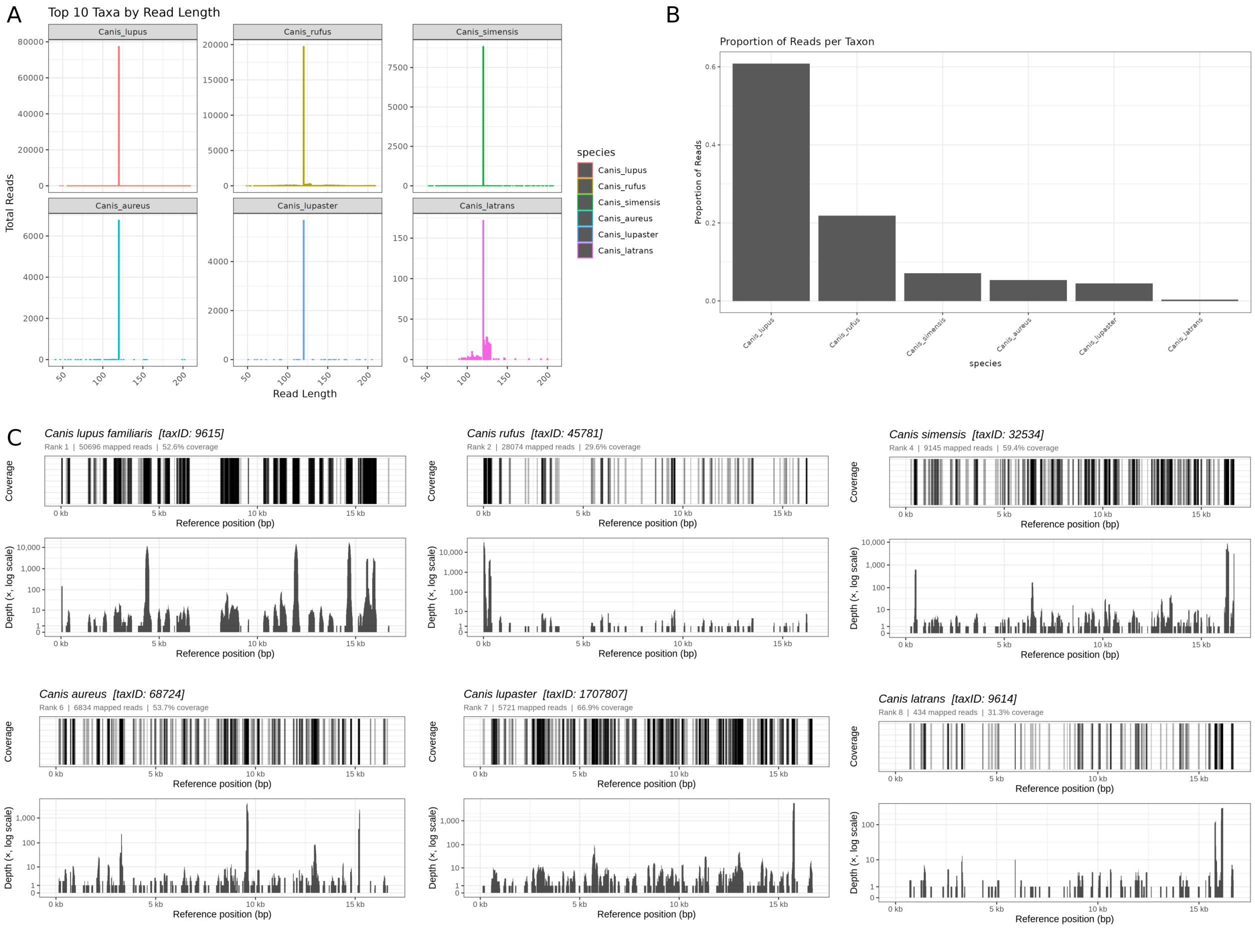


Figure S6: ECMSD output for Clup01 (modern wolf, SRR780933). (A) Read-length distributions of the most abundant classified taxa. (B) Proportion of mitochondrial reads assigned to each canid taxon; *C. lupus* accounts for 60.85% of the 128,249 classified reads. (C) Breadth coverage and per-base depth (log scale) of the *C. lupus* mitochondrial genome (332.9× mean coverage, 52.6% breadth). Full per-sample values are given in the corresponding main-text table.

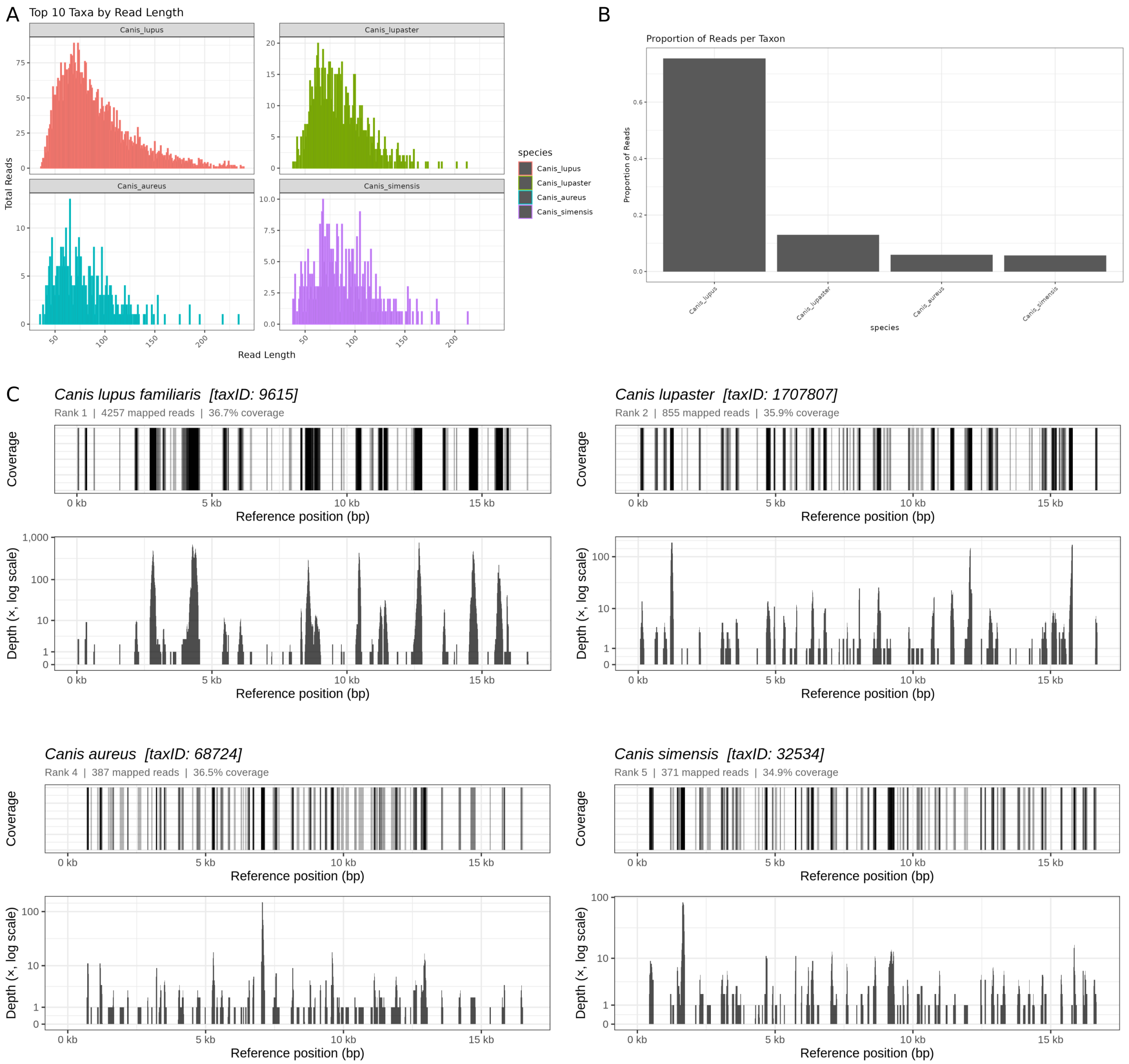


Figure S7: ECMSD output for Clup03 (Neolithic dog, ~7,000y BP, SRR3417117). (A) Read-length distributions of the most abundant classified taxa. (B) Proportion of mitochondrial reads assigned to each canid taxon; *C. lupus* accounts for 75.48% of the 6,579 classified reads. (C) Breadth coverage and per-base depth (log scale) of the *C. lupus* mitochondrial genome (18.7× mean coverage, 36.7% breadth). Full per-sample values are given in the corresponding main-text table.

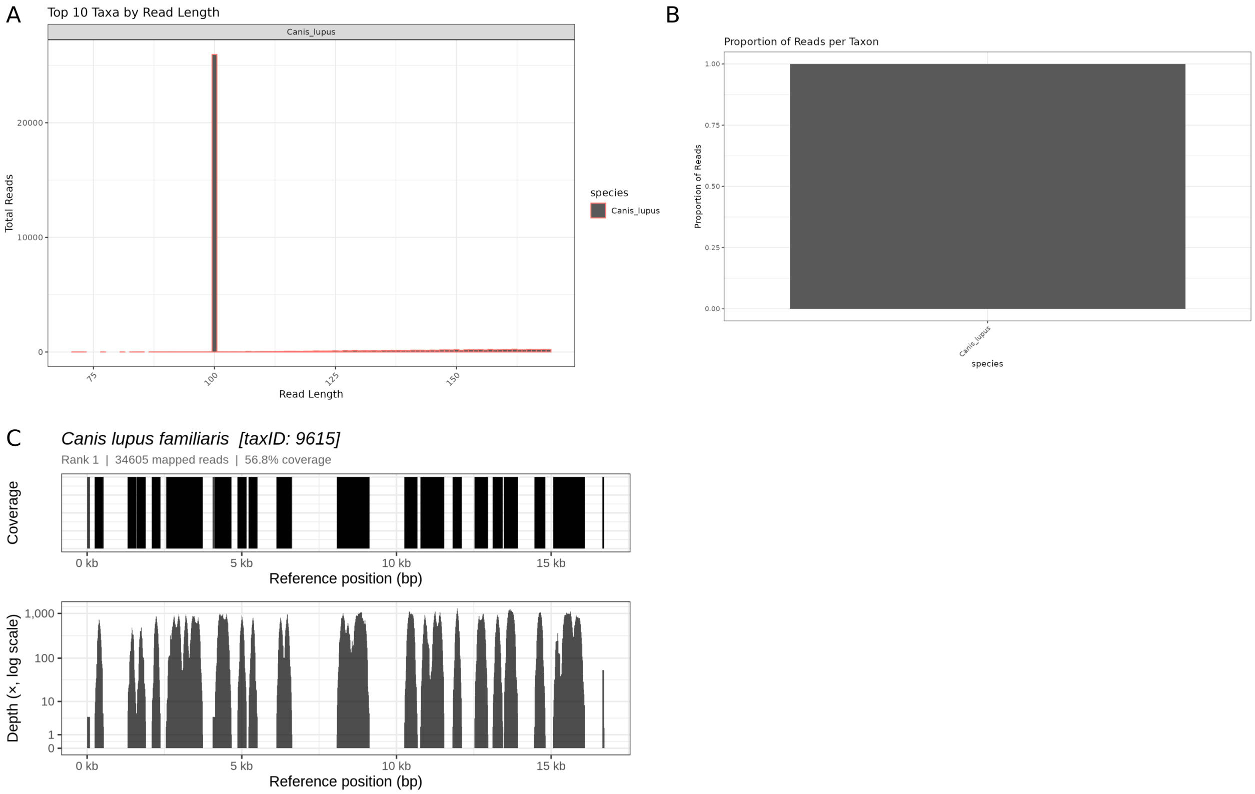


Figure S8: ECMSD output for Clup04 (modern German Shepherd, SRR1122359). (A) Read-length distributions of the most abundant classified taxa. (B) Proportion of mitochondrial reads assigned to each canid taxon; 100% of the 34,605 classified reads were assigned to *C. lupus*. (C) Breadth coverage and per-base depth (log scale) of the *C. lupus* mitochondrial genome (207.2× mean coverage, 56.8% breadth). Full per-sample values are given in the corresponding main-text table.

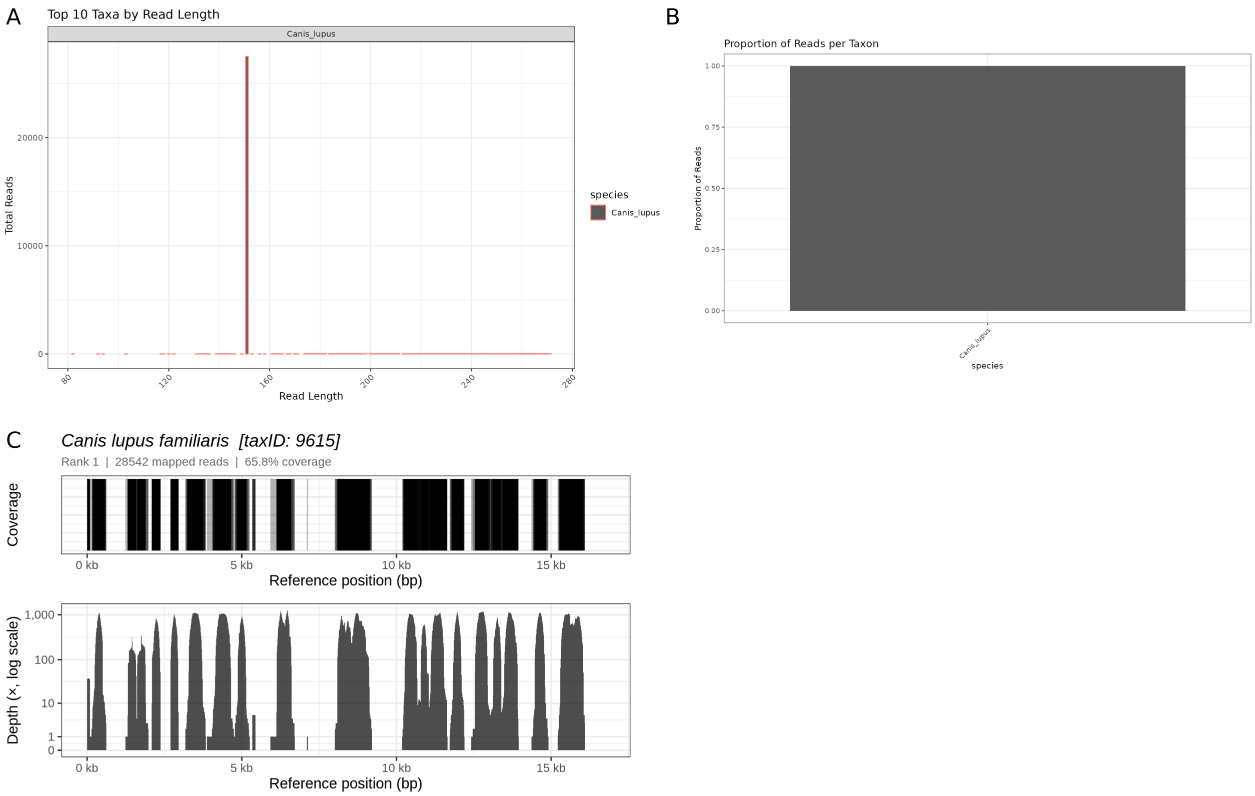


Figure S9: ECMSD output for Clup05 (modern German Shepherd, SRR13339247). (A) Read-length distributions of the most abundant classified taxa. (B) Proportion of mitochondrial reads assigned to each canid taxon; 100% of the 28,542 classified reads were assigned to *C. lupus*. (C) Breadth coverage and per-base depth (log scale) of the *C. lupus* mitochondrial genome (242.4× mean coverage, 65.8% breadth). Full per-sample values are given in the corresponding main-text table.

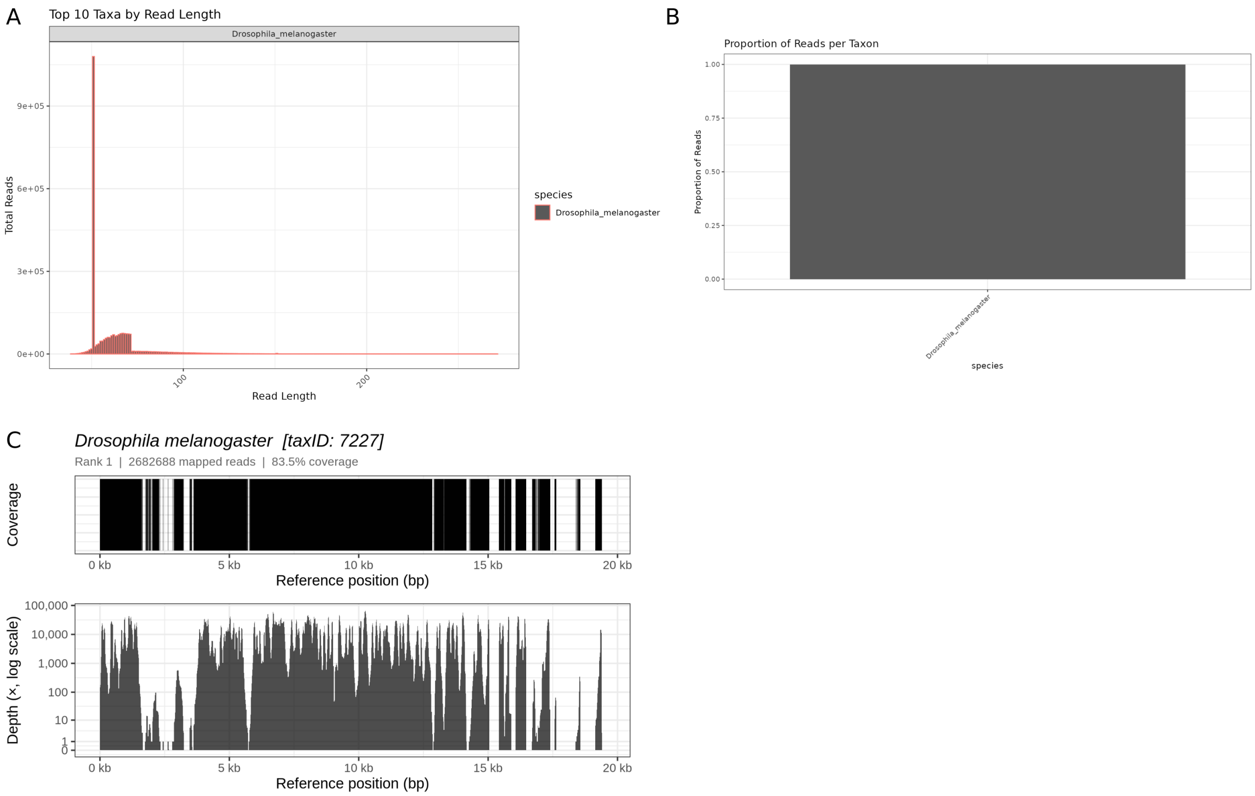


Figure S10: ECMSD output for Dmel1800s-2-GP9 (mid-1800s museum specimen, SRR23876562). (A) Read-length distributions of the most abundant classified taxa. (B) Proportion of mitochondrial reads per taxon; 100% of the 2,682,688 classified reads were assigned to *D. melanogaster*. (C) Breadth coverage and per-base depth (log scale) of the *D. melanogaster* mitochondrial genome (7282.6× mean coverage, 83.5% breadth). Full per-sample values are given in the corresponding main-text table.

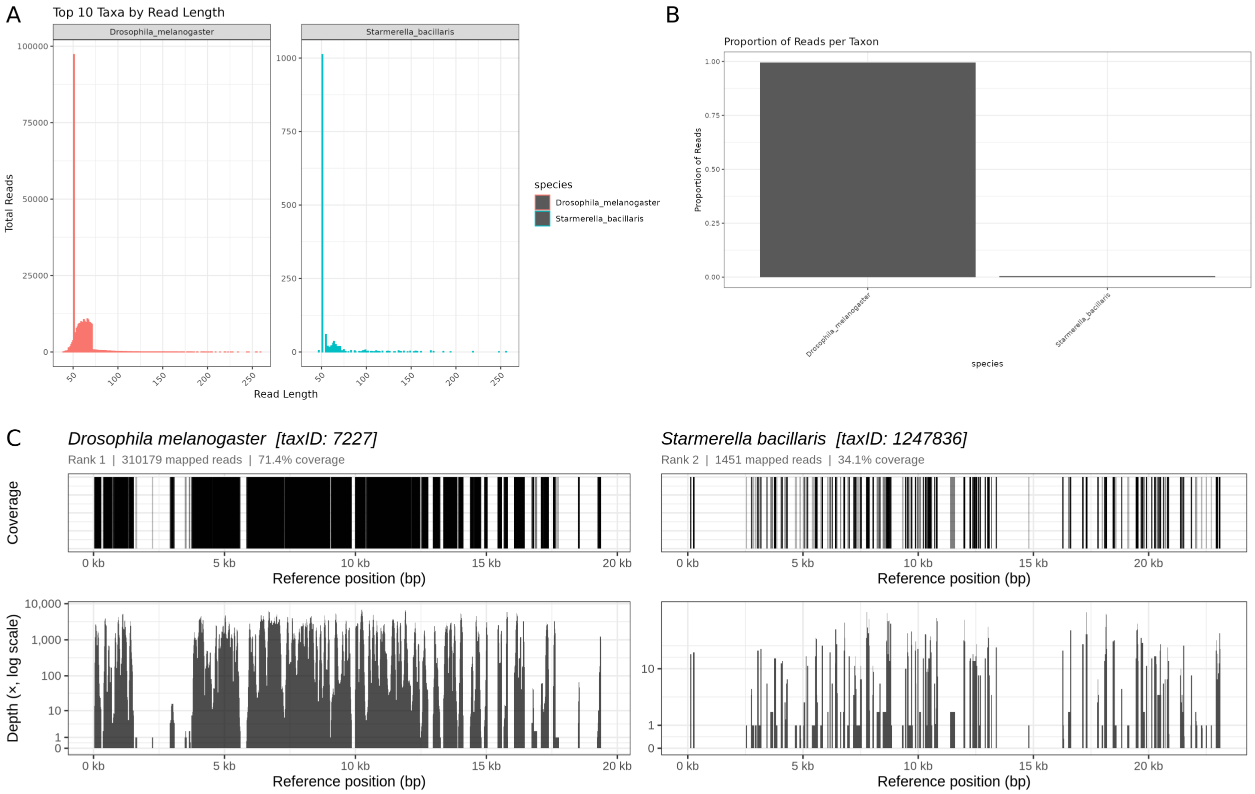


Figure S11: ECMSD output for Dmel1933-SL16 (1933 museum specimen, SRR23876579). (A) Read-length distributions of the most abundant classified taxa. (B) Proportion of mitochondrial reads per taxon; *D. melanogaster* accounts for 99.53% and *Starmerella bacillaris* for 0.47% of the 311,630 classified reads. (C) Breadth coverage and per-base depth (log scale) of the *D. melanogaster* mitochondrial genome (802.9× mean coverage, 71.4% breadth). Full per-sample values are given in the corresponding main-text table.

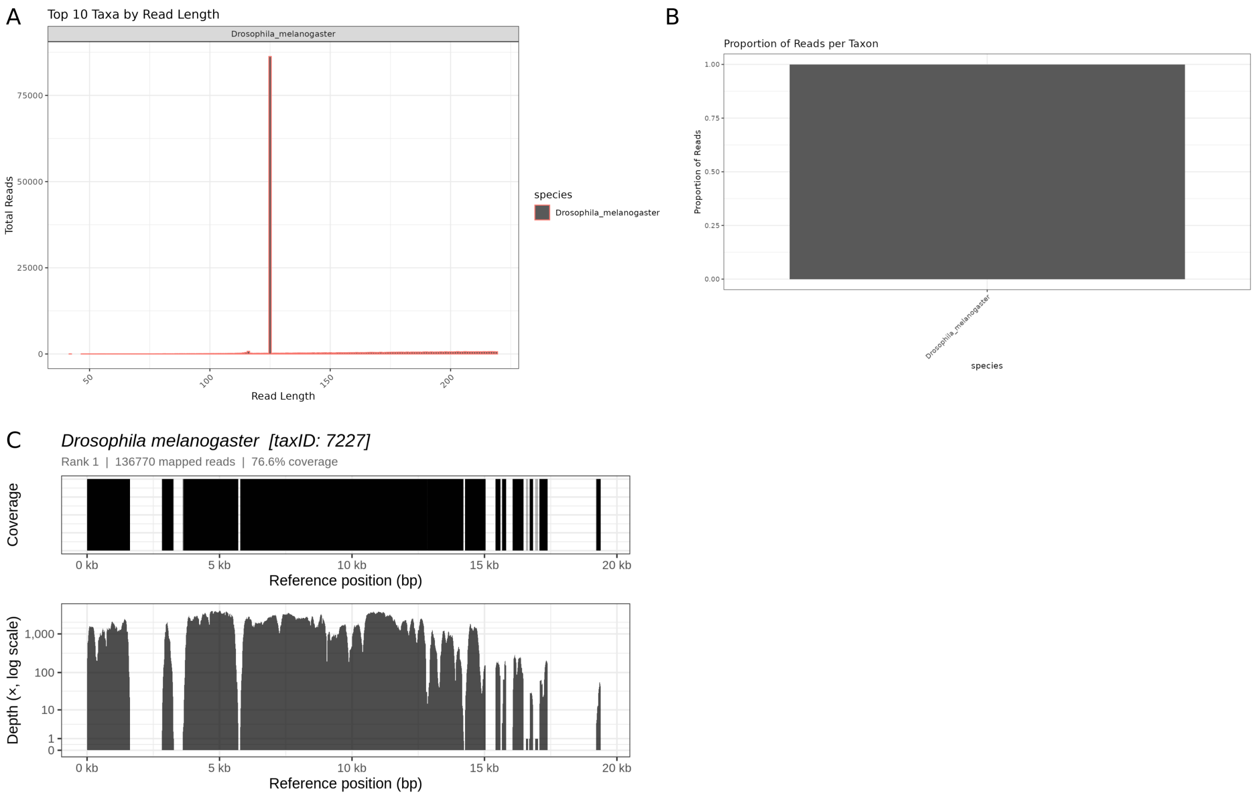


Figure S12: ECMSD output for Dmel1959 (1959 laboratory strain, SRR11846565). (A) Read-length distributions of the most abundant classified taxa. (B) Proportion of mitochondrial reads per taxon; 100% of the 136,770 classified reads were assigned to *D. melanogaster*. (C) Breadth coverage and per-base depth (log scale) of the *D. melanogaster* mitochondrial genome (898.9× mean coverage, 76.6% breadth). Full per-sample values are given in the corresponding main-text table.

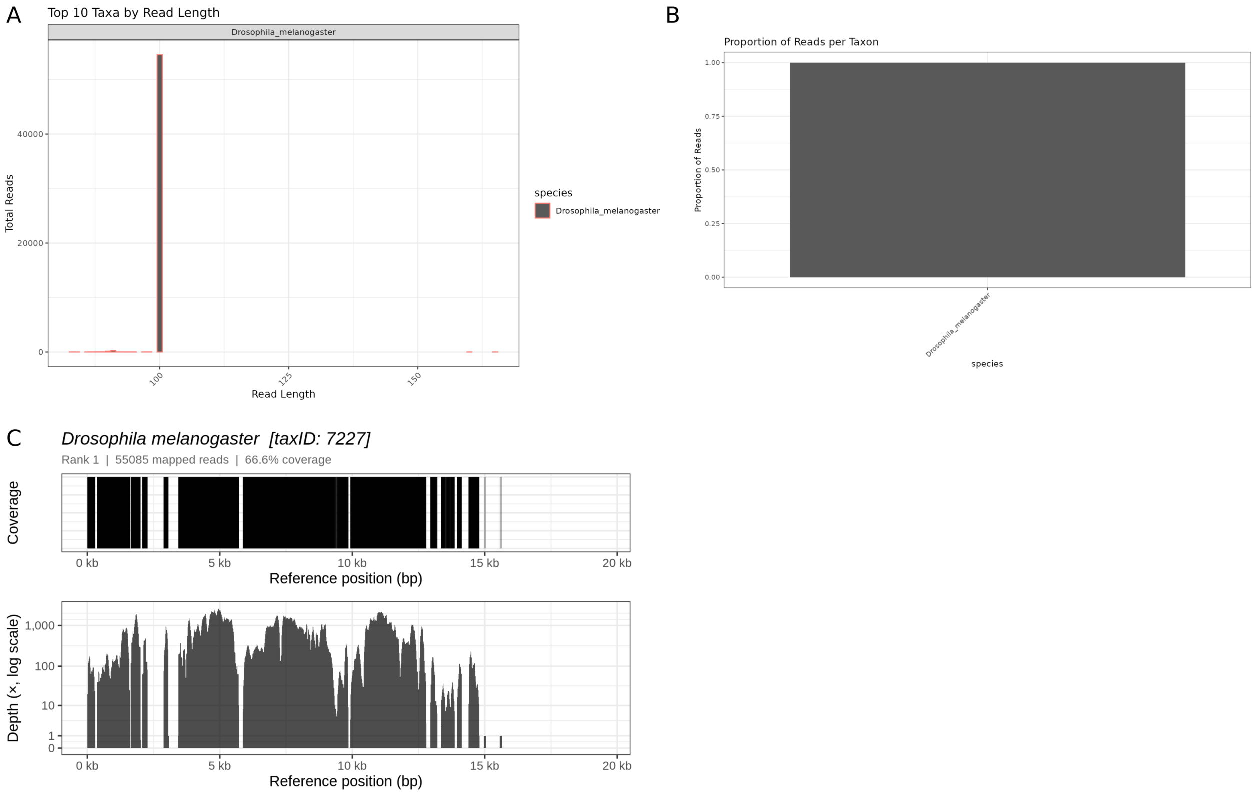


Figure S13: ECMSD output for Dmel2018-STO11 (modern specimen, SRR5762774). (A) Read-length distributions of the most abundant classified taxa. (B) Proportion of mitochondrial reads per taxon; 100% of the 55,085 classified reads were assigned to *D. melanogaster*. (C) Breadth coverage and per-base depth (log scale) of the *D. melanogaster* mitochondrial genome (252.6× mean coverage, 66.6% breadth). Full per-sample values are given in the corresponding main-text table.

### Supplementary Tables

Table S1: Software tools integrated in pastForward v2.0.0. For DeDup we generated a novel fork with increased performance but identical output (see Supplementary Note 1)

| Tool | Version | Function | Reference |
|---|---|---|---|
| Snakemake | 9.9.0 | Workflow management | [Mölder et al., 2021] |
| BUSCO | 6.1.0 | Single-copy gene identification (Module 3) | [Tegenfeldt et al., 2025] |
| ECMSD | 1.3.0 | Mitochondrial taxonomic screen | this study |
| REVEAL | 1.0.0 | Genomic feature dynamics | this study |
| fastp | 1.1.0 | Adapter trimming, QF, read merging | [Chen et al., 2018] |
| BWA-ALN | 0.7.19 | Read alignment (Module 2, Module 3) | [Li and Durbin, 2009] |
| BWA-MEM2 | 2.3 | Read alignment (Module 2, Module 3) | [Vasimuddin et al., 2019] |
| minimap2 | 2.30 | Read alignment (Module 2, Module 3, within ECMSD) | [Li, 2018] |
| BBMap | 39.81 | Read filtering within ECMSD | [Bushnell, 2014] |
| DeDup (fork) | 0.13.0 | Duplicate removal | [Peltzer et al., 2016] |
| mapDamage2 | 2.2.3 | Damage profiling and rescaling | [Jónsson et al., 2013] |
| Centrifuge | 1.0.4.2 | Metagenomic classification | [Kim et al., 2016] |
| SAMtools | 1.23 | BAM processing and statistics | [Li et al., 2009] |
| FastQC | 0.12.1 | Per-sample read QC | [Andrews, 2010] |
| MultiQC | 1.33 | Aggregate QC reports | [Ewels et al., 2016] |
| Qualimap | 2.3 | BAM-level QC | [Okonechnikov et al., 2016] |
| Preseq | 3.2.0 | Library complexity (c_curve) | [Daley and Smith, 2013] |
| pysam | 0.23.3 | BAM/SAM parsing for REVEAL | [Li et al., 2009] |
| R | 4.4.0 | Statistical computing environment | [R Core Team, 2024] |
| ggplot2 | 3.5.1 | Data visualisation | [Wickham, 2016] |

Table S2: Snakemake wrapper versions used in pastForward. All wrappers are from the Snakemake wrappers repository [Mölder et al., 2021]. DeDup, Centrifuge, ECMSD, REVEAL, and Preseq (c_curve) are invoked via direct shell calls as no official Snakemake wrappers exist for these tools.

| **Tool** | **Wrapper path** | **Wrapper version** |
|---|---|---|
| fastp | bio/fastp | v9.3.0 |
| BWA-ALN | bio/bwa/aln, bio/bwa/samse | v9.3.0 |
| BWA-MEM2 | bio/bwa-mem2/mem | v9.3.0 |
| minimap2 | bio/minimap2/aligner | v9.3.0 |
| mapDamage2 | bio/mapdamage2 | v9.3.0 |
| SAMtools | bio/samtools/* | v9.3.0 |
| FastQC | bio/fastqc | v9.3.0 |
| MultiQC | bio/multiqc | v9.3.0 |
| Qualimap | bio/qualimap/bamqc | v9.3.0 |
| BUSCO | bio/busco | v9.3.0 |

Table S3: *C. lupus* samples used in this study (Case Study 1).

| **Sample ID** | **Species** | **Breed** | **Date** | **Origin** | **Accession** | **BioProject** | **Publication** |
|---|---|---|---|---|---|---|---|
| Clup01 | *Canis lupus* | Wolf | Modern | Eurasia | SRR780933 | PRJNA192935 | [Wang et al., 2013] |
| Clup02 | *C. l. familiaris* | Ancient dog | ∼5,000 BP | Germany | SRR3417116 | PRJNA319283 | [Botigué et al., 2017] |
| Clup03 | *C. l. familiaris* | Ancient dog | ∼7,000 BP | Germany | SRR3417117 | PRJNA319283 | [Botigué et al., 2017] |
| Clup04 | *C. l. familiaris* | German Shepherd | Modern | China | SRR1122359 | PRJNA233638 | [Gou et al., 2014] |
| Clup05 | *C. l. familiaris* | German Shepherd | Modern | USA | SRR13339247 | PRJNA685036 | [Evans et al., 2021] |

Table S4: *D. melanogaster* samples used in this study (Case Study 2).

| **Sample ID** | **Species** | **Abbreviation** | **Date** | **Origin** | **Accession** | **BioProject** | **Publication** |
|---|---|---|---|---|---|---|---|
| Dmel1800s-1 | *D. melanogaster* | SL04 | early 1800s | Sweden | SRR23876564 | PRJNA945389 | [Shpak et al., 2023] |
| Dmel1800s-2 | *D. melanogaster* | GP9 | mid 1800s | Germany | SRR23876562 | PRJNA945389 | [Shpak et al., 2023] |
| Dmel1933 | *D. melanogaster* | SL16 | 1933 | Sweden | SRR23876579 | PRJNA945389 | [Shpak et al., 2023] |
| Dmel1959 | *D. melanogaster* | J65 | 1959 | Japan | SRR11846565 | PRJNA634847 | [Schwarz et al., 2021] |
| Dmel2018 | *D. melanogaster* | STO11 | 2018 | Sweden | SRR5762774 | PRJNA390275 | [Mateo et al., 2018] |

### Supplementary Note 1: Accelerated Deduplication (DeDup Performance Fork)

#### Changes

The fork (DeDup-0.13.0) is based on upstream DeDup-0.12.9. It keeps the same command-line interface, output formats, and duplicate detection as the original tool. We made four changes in order to improve performance. $n$ is the local read depth (the number of reads stacked at one locus).

- **Cached per-read fields.** The original re-determines the read-type prefix (`M_` (merged)/`F_` (forward)/`R_` (reverse)) and the summed base quality again for every pairwise comparison. The fork computes both values once, when a read enters the buffer.
- **Hash-index duplicate lookup in merged mode.** In merged mode, all reads are considered merged, whatever their prefix. The read to keep is the one with the highest quality among reads that share the same start and end. The original scans the whole buffer for each comparison. The fork instead looks reads up in a hash index keyed on (start, end). This cuts the cost from $O(n^2)$ to $O(n \log n)$. The BAM output is identical.
- **Coordinate-indexed duplicate lookup in default (prefix-based) mode.** Outside merged mode, the same conditions for determining duplicates is used as in the original. The original algorithm compares reads with each other, even if they could never end up being duplicates (e.g. no common start/end). The fork skips those comparisons. It replaces the original sliding-window buffer with a custom array-based priority queue that also tracks where each read sits in the array. Two hash indices per window, one on start and one on end, cut each candidate set down to the reads that share one coordinate with the anchor. These candidates are then sorted and tested. This reproduces the original's tie-breaking between equal-quality duplicates exactly. Each read is therefore compared only with reads that can actually match, not with the whole window.
- **Parallel per-chromosome processing.** In file mode, each chromosome gets its own worker thread. Each thread reads its own records through the BAM index [Li et al., 2009]. Decompression and record parsing therefore run in parallel. In contrast, the original Dedup processes one chromosome after the other.

#### Verification

The fork's output is byte-identical to the original's. We checked this in three settings: (1) all runtime scenarios, sequential and parallel; (2) synthetic BAMs ranging from 100,000 to 100 million reads, in default and merged mode, and (3) the high-depth pileup scenarios described below. The BAMs matched in all scenarios.

The original JUnit test suite grew from 19 tests to 27. All added tests cover the new functionality and algorithmic changes to ensure correct execution. All 27 tests pass.

#### Performance benchmarks

Table S5: Scaling benchmark, default (prefix-based) mode. Same 15-contig, 300 Mbp synthetic genome at every scale, run on 8 logical CPUs. Values are means of 5 runs per scale.

| Reads | Original | Fork (1T) | Fork (8T) | Speed-up (1T) | Speed-up (8T) |
|---|---|---|---|---|---|
| 100,000 | 1.2 s | 1.1 s | 1.4 s | 1.12× | 0.88× |
| 1,000,000 | 7.1 s | 6.4 s | 6.5 s | 1.12× | 1.10× |
| 5,000,000 | 22.1 s | 16.6 s | 14.7 s | 1.33× | 1.51× |
| 10,000,000 | 43.8 s | 25.1 s | 16.1 s | 1.77× | 2.74× |
| 25,000,000 | 95.2 s | 46.1 s | 20.5 s | 2.07× | 4.68× |
| 50,000,000 | 210.1 s | 83.8 s | 26.1 s | 2.50× | 8.06× |
| 100,000,000 | 401.7 s | 145.7 s | 34.3 s | 2.78× | 11.84× |

T = threads.

Table S6: Scaling benchmark, merged mode. Same 15-contig, 300 Mbp synthetic genome at every scale, run on 8 logical CPUs. Values are means of 5 runs per scale.

| Reads | Original | Fork (1T) | Fork (8T) | Speed-up (1T) | Speed-up (8T) |
|---|---|---|---|---|---|
| 100,000 | 1.1 s | 1.0 s | 1.3 s | 1.10× | 0.82× |
| 1,000,000 | 6.3 s | 5.5 s | 6.3 s | 1.16× | 1.01× |
| 5,000,000 | 22.7 s | 15.8 s | 13.6 s | 1.44× | 1.67× |
| 10,000,000 | 35.6 s | 20.1 s | 14.4 s | 1.78× | 2.47× |
| 25,000,000 | 84.7 s | 37.0 s | 19.1 s | 2.28× | 4.42× |
| 50,000,000 | 166.7 s | 59.4 s | 23.9 s | 2.79× | 6.94× |
| 100,000,000 | 292.8 s | 100.3 s | 31.1 s | 2.92× | 9.61× |

T = threads.

Below roughly 1 million reads, multi-thread mode is typically slower than running as single thread, due to internal thread management. Parallel mode pulls ahead with increasing number of reads and chromosomes. At 100 million reads it is 11.8× (default mode) and 9.6× (merged mode) faster than.

Two further benchmarks target the $O(n^2)$ high-depth pileup case directly. Both use a single contig and run single-threaded. Their speed-ups therefore come from the coordinate-indexed lookup alone, not from the parallel per-chromosome processing. Each scenario ran 5 times on the same BAM. Values shown are means.

Table S7: High-depth local-pileup benchmark, merged scenario (`-m`, BAM generated by `generate_big_pileup_bam.py`): original vs. fork at a single locus. All reads share one start position. **Groups** is the number of distinct (start, end) pairs, i.e. distinct fragment lengths, each with 40 duplicate reads.

| Groups | Reads | Original (mean) | Fork (mean) | Speed-up |
|---|---|---|---|---|
| 500 | 20,000 | 2.0 s | 0.5 s | 4.22× |
| 1,000 | 40,000 | 4.7 s | 0.6 s | 7.46× |
| 2,000 | 80,000 | 17.3 s | 1.0 s | 17.75× |
| 4,000 | 160,000 | 71.2 s | 2.5 s | 28.18× |
| 8,000 | 320,000 | 306.2 s | 7.9 s | 38.80× |

Table S8: High-depth local-pileup benchmark, default scenario (no `-m`, BAM generated by `generate_local_peak_bam.py`): original vs. fork across a narrow band of staggered start positions. **Groups** is the number of distinct start positions, 1 bp apart, each with 10 duplicate reads. Reads are 400 bp long, so the whole band stays buffered in the window at once.

| **Groups** | **Reads** | **Original (mean)** | **Fork (mean)** | **Speed-up** |
|---:|---:|---:|---:|---:|
| 500 | 5,000 | 1.6 s | 0.4 s | 4.54× |
| 1,000 | 10,000 | 4.2 s | 0.6 s | 7.53× |
| 2,000 | 20,000 | 14.0 s | 0.8 s | 16.87× |
| 4,000 | 40,000 | 59.2 s | 1.9 s | 31.27× |
| 8,000 | 80,000 | 273.8 s | 6.0 s | 45.94× |

Speed-up factor increases with depth in both scenarios. The original tool's scan cost per read comparison grows with $n$. The fork's indexed lookups stay close to constant. At the largest tested group count (8,000 groups), the fork is 45.9× faster in default mode and 38.8× faster in merged mode.

#### Availability

The fork is available at `https://github.com/SarahSaadain/DeDup`. It was forked from the original DeDup tool at `https://github.com/apeltzer/DeDup` [Peltzer et al., 2016]. The benchmarking and correctness comparison used to generate the results above is available at `https://github.com/SarahSaadain/DeDup_comparison_fork`.

#### Supplementary Data

**Supplementary Data 1.** Full MultiQC summary report (PDF) generated by the Summary Module for the *C. lupus* case study, aggregating read, mapping, and coverage statistics across all five samples.

**Supplementary Data 2.** Full MultiQC summary report (PDF) generated by the Summary Module for the *D. melanogaster* case study, aggregating read, mapping, and coverage statistics across all samples.
